# RNA 3D Motif Dynamics Guide Assembly of the Replication Initiation Complex in Flaviviruses

**DOI:** 10.64898/2026.06.16.732768

**Authors:** Lorena V. Streit, Takashi Onikubo, Miro A. Astore, Emily A. Cioppa, Paul Dominic B. Olinares, Linas Urnavicius, Danny Incarnato, Steve L. Bonilla

## Abstract

Viral RNA genomes contain structured elements that recruit the machinery necessary for their replication. However, these mechanisms remain poorly understood because of RNA structure’s dynamic nature. Using cryo-EM and single-molecule Förster resonance energy transfer, we show that the flaviviral stem-loop A (SLA)—which recruits the viral polymerase NS5 to initiate negative-strand synthesis—populates a conserved, multi-state conformational ensemble governed by a dynamic three-dimensional (3D) motif. Structures of Zika and dengue SLA–NS5 complexes reveal that NS5 engages a preorganized SLA conformation. Notably, destabilizing this conformation impairs negative-strand synthesis without affecting NS5 binding. Thus, the conserved SLA 3D ensemble guides a post-binding conformational search that converts an initial encounter complex into a productive replication-initiation complex—revealing how RNA conformational ensembles, rather than static structures, dictate function.

**SUMMARY:** Viral genomes encode dynamic RNA structures that are specifically recognized by proteins to regulate critical steps of the viral life cycle. Although these RNAs exist as multi-state conformational ensembles, structural studies have largely captured single conformations, limiting our mechanistic understanding of RNA-protein recognition and function. Here, using single-particle cryo-electron microscopy (cryo-EM), single-molecule Förster resonance energy transfer, and biochemical assays, we show how flaviviral genomes use structural dynamics encoded in their stem-loop A (SLA) to guide a conserved, multi-step mechanism of negative-strand synthesis initiation by non-structural protein 5 (NS5). Cryo-EM ensembles of SLAs from dengue, Zika, West Nile, and yellow fever viruses reveal a conserved conformational landscape dictated by alternative stacking configurations of a central three-way junction (3WJ). Cryo-EM structures of SLA-NS5 complexes from dengue and Zika show that NS5 engages a specific state within SLA 3D ensemble, one in which the 3WJ adopts a GRR/A tetraloop-like motif that we identify across diverse structural RNAs. Strikingly, mutations that substantially destabilize this state impair negative-strand synthesis without affecting NS5 binding—decoupling binding from function and ruling out a simple conformational-capture mechanism. Instead, the SLA’s intrinsic structural dynamics direct a post-binding conformational search that converts an initial, conformationally heterogeneous encounter complex into a productive replication-initiation complex. These results demonstrate a mechanism of RNA-protein recognition in which the 3D conformational ensemble of an RNA guides formation of the functional state downstream of binding, and identify a ubiquitous, intrinsically dynamic 3D motif within the SLA’s 3WJ as a critical determinant of flavivirus negative-strand synthesis.

## INTRODUCTION

Viruses encode functional RNA three-dimensional (3D) structures that act in concert with viral and host proteins to initiate and regulate critical viral processes (*1–3*). Yet, despite their importance and therapeutic potential, our mechanistic understanding of how RNA structures regulate these processes remains limited. A major challenge lies in their dynamic nature, as these RNA elements do not exist as single, static structures but instead populate dynamic conformational ensembles that are difficult to capture experimentally (*4–7*). These ensembles can include conformations that engage with and regulate the function of their binding partners, as well as alternative conformations that are not directly recognized yet shape the pathways leading to functional states (*4, 8, 9*). Therefore, a mechanistic understanding of functional viral RNA elements requires dissection of their dynamic 3D conformational ensembles.

Flaviviruses are positive-sense single-stranded RNA (+ssRNA) viruses that include important human pathogens such as dengue virus (DENV), Zika virus (ZIKV), West Nile virus (WNV), and yellow fever virus (YFV) (*10*). DENV alone is estimated to infect ∼400 million people each year, and global changes in climate, urbanization, and travel are expected to further increase the incidence of flaviviral infections worldwide (*10–12*). Despite this global burden, there are no approved antivirals for flaviviruses, underscoring the need to identify mechanisms that can be therapeutically targeted.

Like other +ssRNA viruses, flaviviruses rely on RNA structures to regulate critical steps of their life cycles (*13–15*). A conserved ∼70-nucleotide RNA element known as stem-loop A (SLA) is encoded in the 5′ end of the ∼11 kilobase genome and is required for genome replication by interacting with non-structural protein 5 (NS5) to initiate synthesis of a negative-strand using the +ssRNA as a template (*15–17*). NS5 is the largest and most conserved flaviviral protein and comprises an RNA-dependent RNA polymerase (RdRp) domain that catalyzes *de novo* RNA synthesis and a methyltransferase (MTase) domain that harbors enzymatic activities required for the formation of a type-1 cap structure at the 5′ end of the viral genome (*18–20*). Negative-strand synthesis initiates at the 3′ end of the genome, a process that requires long-range RNA-RNA interactions that circularize the genome and bring the 3′ end in proximity to SLA (*21–25*). Whether SLA serves as a passive binding site for NS5 recruitment or whether its structural dynamics play an active regulatory role in NS5 recognition and negative-strand synthesis is not understood.

SLAs from different flaviviruses can vary in sequence, size, and detailed secondary structural features (e.g., 2-way junctions, bulges, etc.) (*26*). Despite these differences, bioinformatic, chemical probing, and NMR studies consistently reveal a conserved global secondary structure in which bottom (B), top (T) and side (S) stems join at a central three-way junction (3WJ) to form a “Y-shaped” architecture (Fig. 1A) (*17, 26–30*). The strong conservation of NS5 and the shared secondary structure across SLA variants support a conserved mechanism of SLA-NS5 recognition. Consistent with this, SLAs from diverse flaviviruses can substitute for one another (*31–35*). Yet experimental static 3D structures of SLAs from different flaviviruses have produced divergent conformations, leading to models of species-specific SLA-NS5 recognition modes (*31, 35, 36*). These structures may instead represent distinct states within a conserved, dynamic ensemble. To reconcile the apparent conflict between conserved function and divergent static structures, and to reveal how SLA structural dynamics regulate NS5 function, we leveraged single-particle cryo-electron microscopy (cryo-EM) and single-molecule Förster resonance energy transfer (smFRET) to dissect the 3D ensembles of SLAs from diverse flaviviruses in isolation and in complex with NS5.

**Fig. 1.**
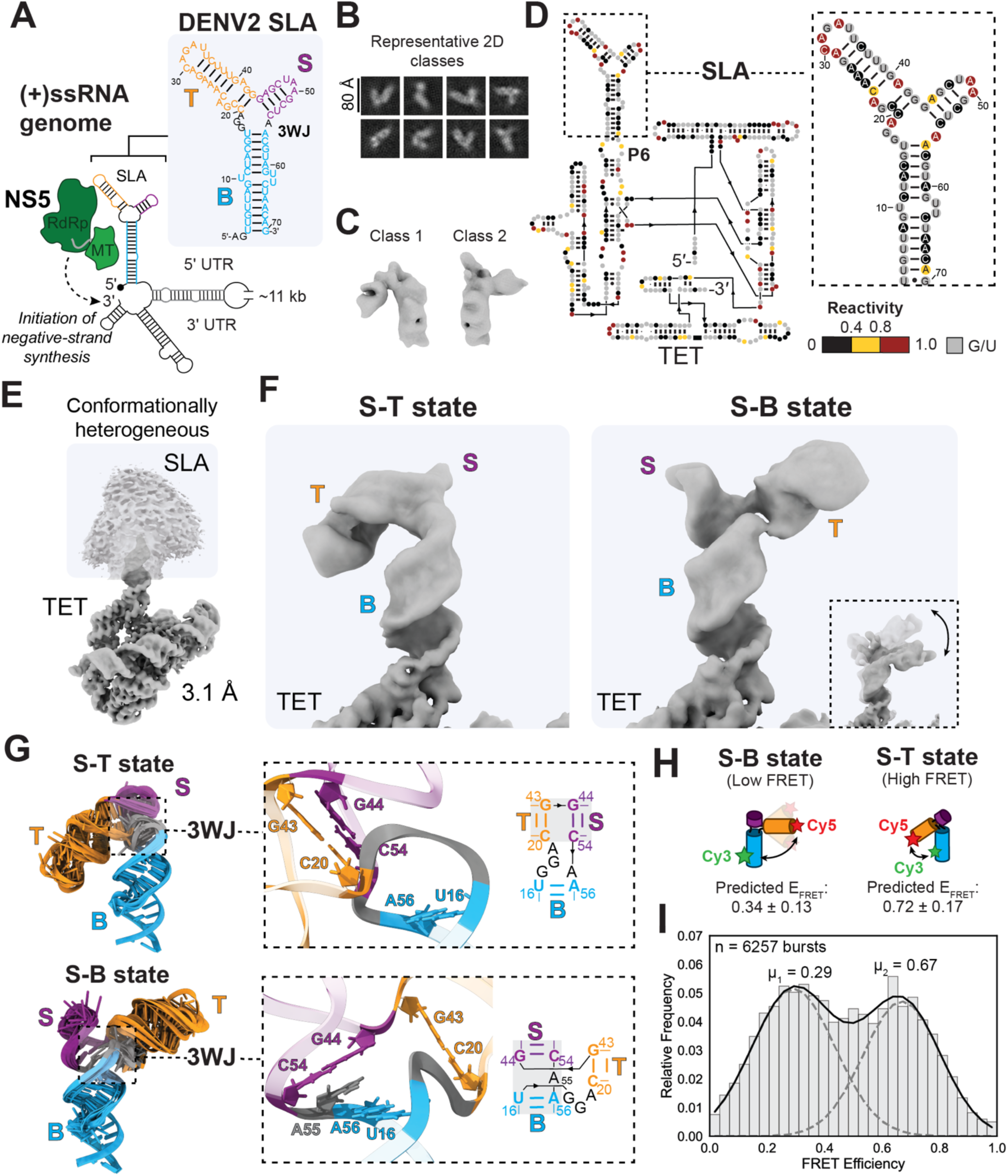
DENV2 SLA adopts multiple 3D conformations governed by the conformational preferences of its 3WJ. (**A**) Sequence and secondary structure of DENV2 SLA in the context of the circularized (+)ssRNA genome. SLA is encoded in the 5′ UTR of the 5′ capped flavivirus genome and comprises bottom (B), top (T), and side (S) stems that join at a three-way junction (3WJ). NS5 recruitment to SLA and genome cyclization facilitates initiation of negative-strand synthesis from the 3′ end. (**B**) Representative cryo-EM 2D projections of DENV2 SLA 70-nt construct. (**C**) Cryo-EM 3D reconstructions of two different conformations of DENV2 SLA 70-nt construct. (**D**) DMS chemical probing of DENV2 SLA appended to the P6 helix of the TET scaffold. (**E**) Refined map of scaffolded DENV2 SLA prior to 3D classification. (**F**) Refined maps of major 3D conformations of scaffolded DENV2 SLA. Densities corresponding to TET, B, T, and S stems are labeled. Additional subclasses of the S-B state are shown in the bottom right corner. (**G**) Top-ten scoring models generated by auto-DRRAFTER of the S-T (top) and S-B (bottom) states. The 3WJ geometry of the median-scored model is shown on the right, with diagrams illustrating the stem-stacking arrangement indicated by grey boxes. (**H**) FRET states predicted for Cy3/Cy5 labeled DENV2 SLA. (**I**) FRET efficiency histogram of DENV2 SLA at 10 mM MgCl_2_ fitted to a two-state model. Normalized FRET efficiency distribution is shown with 30 bins.

Using scaffold-based cryo-EM, we show that SLA variants from DENV, ZIKV, YFV, and WNV are highly structurally dynamic, but that the major 3D conformations are conserved and dictated by shared properties of the 3WJ. Cryo-EM structures of SLA-NS5 complexes from ZIKV and DENV reveal that NS5 engages a specific conformation within the unbound SLA ensemble, one in which S and T stems coaxially stack and the 3WJ adopts a ubiquitous tetraloop (TL)-like 3D motif. We refer to this conformation in the SLA ensemble as the S-T state. The presence of the S-T state and its engagement by NS5 is shared across the different flaviviruses studied, supporting a conserved mechanism of recognition. Mutations that destabilize the TL-like motif of the S-T state severely perturb the SLA 3D ensemble and impair negative-strand synthesis but, strikingly, they do not affect NS5 binding. This supports a mechanism in which the intrinsic structural dynamics of SLA guide a post-binding conformational search for the replication initiation-competent complex. These findings demonstrate how RNA 3D structural dynamics can affect RNA-protein recognition and function beyond initial binding and highlight the 3WJ of SLA as a potential therapeutic target.

## RESULTS

### SLA forms multiple, globally distinct 3D conformations

We used single-particle cryo-EM to visualize SLA elements from different flaviviruses, beginning with a minimal construct encoding the 70 nt SLA from DENV serotype 2 (DENV2; Fig. 1A). Despite the small size of this construct (∼23 kDa), cryo-EM analysis produced 2D classes and 3D reconstructions consistent with the size and expected Y-shaped secondary structure of DENV2 SLA (Fig. 1B&C). Notably, unlike prior studies reporting single static structures, our initial cryo-EM analysis suggested that SLA adopts multiple conformations (Fig. 1C). However, due to the small size of the construct and the limited resolution of these maps, the relative orientation of the helices could not be inferred unambiguously, and the distinct 3D conformations could not be determined.

To overcome this, we employed a scaffold-based approach that leverages the well-established modular nature of RNA structure, improves cryo-EM particle alignment, and facilitates assignment of secondary structural domains for 3D modelling (*37, 38*). We used the *Tetrahymena* ribozyme (TET) as a scaffold, as it has been shown to serve as a robust scaffold that does not interfere with the native fold of appended RNAs and its structure has been determined to high resolution by cryo-EM (*37, 39*). Using this approach, we generated RNA constructs where the B stem of the DENV2 SLA is appended to P6 of TET, effectively replacing TET’s P6b stem (Fig. 1D). Dimethyl sulfate mutational profiling with sequencing (DMS-MaPseq) confirmed that the previously determined secondary structures of DENV2 SLA and TET are maintained in the scaffold construct (Fig. 1D; fig. S1).

Cryo-EM analysis produced a map with a global resolution of ∼3.1 Å (Fig. 1E; fig. S2). The density corresponding to TET showed excellent agreement with its previously determined atomic structure (fig. S2), further supporting that TET and SLA fold into their native structures. Consistent with our initial results with the 70-nt construct (Fig. 1C), the density corresponding to the DENV2 SLA displayed conformational heterogeneity (Fig. 1E).

To characterize the SLA’s conformational ensemble—rather than focusing only on the dominant state, as is typical in cryo-EM workflows—we examined the distribution of particles across all 3D classes (Fig. 1F; fig. S2 & S3). Analysis of the conformational heterogeneity of SLA using focused 3D classification revealed two dominant conformational states that differed in how the B, T, and S stems are oriented relative to each other (Fig. 1F). One of these states (S-T state) was well represented by a single 3D class, while the other one (S-B state) was more flexible, suggesting the presence of substates (Fig. 1F). The S-T and S-B states were robustly reproduced when different numbers of classes were used for 3D classification (fig. S3). The large differences in the relative orientations of the B, T, and S stems suggest substantial conformational changes between these states. Their shapes qualitatively matched those observed with the 70-nt SLA construct (fig. S4).

Our semi-quantitative analysis indicated that ∼40-55% of the single particles occupy the S-T and S-B states, while other particles were assigned to classes that did not produce well-defined 3D reconstructions (fig. S3). This suggests the presence of additional, dynamic and broadly distributed states. These states likely include intermediates in a continuum between the S-T and S-B states and may also include other conformations, such as those with a melted S stem, consistent with local dynamics observed by NMR studies (*29*). Notably, we found a small fraction of the total particles (∼6–10%) forming dimers that agreed with those observed in a previously reported crystal structure of dimerized DENV2 SLAs (*35*), in which the canonical Y-shaped secondary structure is disrupted by formation of intermolecular base pairs between melted S stems (fig. S2). Although these dimers are unlikely to represent a functional state—as they require high RNA concentrations to form (*28*) and are not observed in SLA-NS5 complexes described below—their presence is consistent with the SLA sampling conformations in which the S stem melts. In the crystal structures, these conformations would have been stabilized by formation of intermolecular base pairs, facilitated by the high RNA concentrations in the crystallization conditions.

To test whether the unresolved states observed in cryo-EM include conformations in which the S stem is melted, we repeated our cryo-EM studies with a construct designed to stabilize the S stem. Using a previously validated circularly permuted TET (cpTET) scaffold (*37*), we produced a construct in which the S stem of DENV2 SLA—rather than the B stem—is appended to P6, stabilizing the S stem (fig. S5). Focused 3D classification produced the same S-T and S-B states observed in the original scaffold, confirming that these states are intrinsic to the SLA and independent of scaffold context (fig. S5 & S6). Although some unresolved classes still emerged, their fraction was substantially reduced and ∼80% of particles classified into the S-T and S-B states (fig. S5). Stabilizing the S stem thus shifted the population from unresolved states toward the S-T and S-B states, supporting the conclusion that the dynamic, broadly distributed states in the ensemble include conformations with a melted S stem. Consistent with S stem stability playing a role on the conformational properties of the SLA, its perturbation leads to replication defects *in vitro* and *in vivo* (*15, 17*).

Overall, our 3D reconstructions and semi-quantitative analysis show that DENV2 SLA populates multiple 3D conformations: the S-T and S-B states are sufficiently stable to be resolved by cryo-EM, while the remaining particles occupy a continuum of dynamic states—including conformations with a melted S stem—whose relative populations are tuned by S stem stability.

### Alternative stacking configurations of 3WJ dictate the global 3D ensemble of SLA

Despite the moderate local resolution of the maps at the SLA region, the global 3D architectures of the S-T and S-B states were discernible, and their backbones could be reliably traced. The cryo-EM maps, along with the secondary structure of DENV2 SLA and atomic structure of TET (*39*), were used as restraints for Rosetta-based automated 3D modeling, implemented in auto-DRRAFTER (Fig. 1G) (*40*). Auto-DRRAFTER fits multiple 3D models into the density; we used the top ten models ranked by its internal scoring to represent modeling uncertainty, as done previously (*41, 42*).

While the conformation of bases at non-canonical regions could not be determined with atomic precision due to the moderate resolution of the maps, the base-paired regions and the backbone conformations—including at the critical 3WJs—are consistent across 3D models (Fig. 1G; fig. S2). Comparison of the S-T and S-B states reveals substantial differences in the internal conformations of their 3WJs, with different stacking arrangements of the B, T, and S stems, producing large differences in the relative orientations of the emanating stems (Fig. 1G; fig. S7). The 3D models indicate that the heterogeneity observed in the S-B state is well represented by substates that share the same helical stacking arrangement but differ in the angle between the B and T stems (fig. S2); for simplicity, we use a single representative S-B conformation for comparisons hereafter. The S-T state, by contrast, is more rigid and is well-represented by a single conformation.

In the S-T state, the S and T stems coaxially stack via their closing base pairs, C20-G43 in the T stem and G44-C54 in the S stem, requiring a sharp turn of the strand containing residues G17, G18, and A19 in the 3WJ (Fig. 1G, top). The S-T coaxial stacking is consistent with the overall architecture proposed for DENV1 SLA by NMR (*28*), although the two models differ in the internal configuration of the 3WJ and in the relative orientation of the B stem—possibly reflecting contributions from multiple conformations to the NMR data that could not be resolved as discrete states. In the S-B state, the S and B stems stack to form a semi-continuous helix facilitated by A55 of the 3WJ, which stacks with C54 and A56 to “bridge” the S and B stems, as seen consistently in all 3D models of the S-B state (Fig. 1G, bottom). These alternative stacking configurations result in a ∼180° difference in the orientations of the B and T stems when comparing the S-T and S-B states (fig. S7). Additionally, the states differ in their compactness. In the S-B state, there is a ∼110° angle between the B and T stems resulting in an extended structure, while in the S-T state this angle is ∼60°, resulting in a more compact structure (fig. S7). Thus, alternative local configurations of the 3WJ produce dramatically different global 3D conformations of SLA.

### smFRET reveals a tunable conformational ensemble in solution

To validate the S-T and S-B states in solution and obtain quantitative information about their relative populations, we generated Cy3/Cy5-labeled constructs for smFRET, with predicted low and high FRET states for the S-B and S-T states, respectively, based on our cryo-EM structures (Fig. 1H; fig. S8). After confirming that labeling does not perturb SLA folding (fig. S8), confocal smFRET measurements yielded a bimodal distribution well described by a two-state model, with mean FRET efficiencies consistent with those predicted from the S-T and S-B structures (Fig. 1H & I; fig. S8; Table S1). The fluorophores were positioned to report on the global rearrangements that distinguish the S-T and S-B states. Other conformations within the ensemble—including intermediates and S-stem-melted conformations—likely contribute to the observed FRET distribution but are not resolved as additional peaks, possibly because they are too dynamic or sparsely populated to be detected as discrete states and/or because their FRET values fall within the range spanned by the two major peaks. Within this two-state framework, both FRET states are observed across all solution conditions tested, but their relative populations changes with [Mg^2+^] (fig. S8). At low [Mg^2+^], the FRET state corresponding to the S-B state is more populated; the relative population of the S-T state grows with increasing [Mg^2+^] (fig. S8), consistent with its more compact structure and the greater electrostatic neutralization required to counteract RNA-RNA repulsion. Together with a mutation described below that destabilizes the S-T state and shifts the FRET distribution accordingly, these results support a model in which the S-T and S-B states are populated in solution and their relative occupancies are tuned by environmental conditions including [Mg^2+^].

### SLA 3D structural ensemble is conserved

Having established that the global 3D ensemble of DENV2 SLA is governed by alternative stacking configurations of the 3WJ, we next asked whether this property is conserved across flaviviruses. In previous studies, inspection of the predicted secondary structures of SLAs from different flaviviruses suggested that certain properties of the 3WJ—such as the relative lengths of their strands—may be conserved, pointing to a shared 3WJ 3D structure (*28, 33*). To more broadly explore shared 3WJ properties that may indicate conserved conformational dynamics of the SLA, we first used a bioinformatic approach (*43, 44*) to generate a secondary structure covariation-based alignment of SLAs from 35 diverse flavivirus variants, including important mosquito-borne and tick-borne human pathogens (Fig. 2A; fig. S9) Our analysis showed that—in addition to previously reported sequence motifs such as the U-bulge in the B stem and an AG dinucleotide in the apical loop of the T stem (*17, 33, 35, 45*)—several features of the 3WJ are strongly conserved (Fig. 2B).

**Fig. 2.**
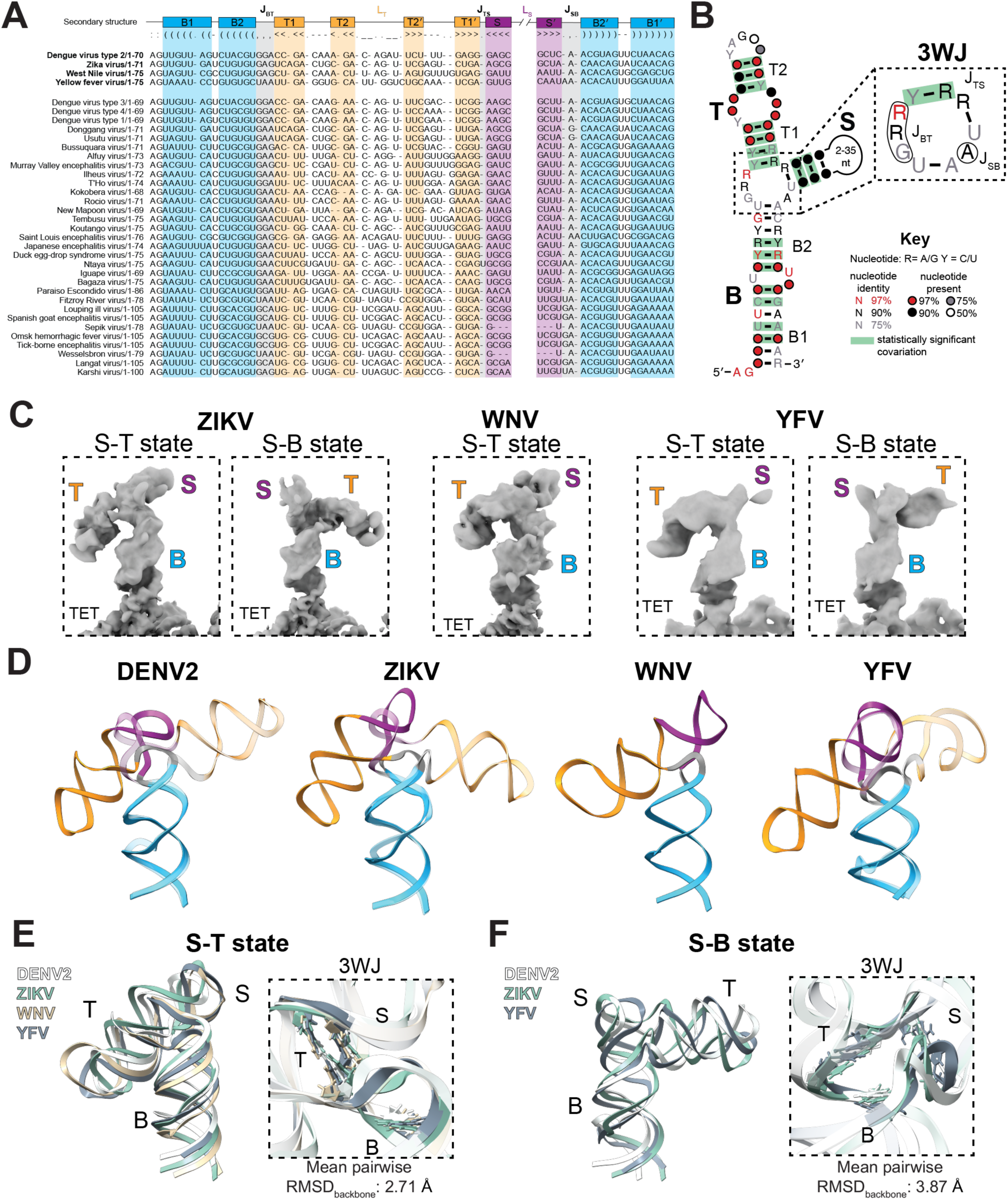
Conservation of SLA 3D ensembles. (**A**) Infernal alignment of SLAs from 35 diverse flaviviruses. The consensus secondary structure and sequences corresponding to SLAs from DENV2, ZIKV, WNV and YFV studied here are shown at the top. Regions corresponding to B (B1 and B2), T (T1 and T2), and S stems are labeled using same colors scheme as in Fig. 1A. S stem insertions are not shown; see fig. S9 for full alignment. (**B**) Covariance models of flavivirus SLA reveals conserved features of the central 3WJ. Covarying base pairs validated with R-scape with E- value < 0.05 are shown in green. The model was generated using the Infernal alignment in panel A. (**C**) Refined cryo-EM maps of major 3D conformations of ZIKV, WNV and YFV SLAs. Regions corresponding to TET, B, T, and S stems are labeled. (**D**) Structures fitted into cryo-EM maps using auto-DRRAFTER (*41*). The S-T and S-B states are superimposed by aligning their B stem. The S-B state is shown in transparent. B, T and S stem are colored using the same color scheme as in panel A. (**E**) Comparison of S-T states across flavivirus SLAs. Median-scored models from auto-DRRAFTER were aligned using global backbone (left) and 3WJ closing base-pair residues (right). Mean pairwise RMSD was calculated using the RNA backbone atoms of the 3WJ including the closing base-pairs (A59 of YFV SLA was excluded for residue matching). (**F**) Comparison of modeled S-B states across flavivirus SLAs. Median-scored models from auto-DRRAFTER were aligned using global backbone (left) and 3WJ closing base-pair residues (right). Mean pairwise RMSD was calculated using the RNA backbone atoms of the 3WJ including the closing base-pairs (A59 of YFV SLA was excluded).

First, there is strong conservation of a G and an A at the base of the 3WJ (e.g., G17 and A55 in DENV2). Second, almost invariably, the T and the S stems are directly connected without intervening residues. In addition, the junction connecting the S and B stems (J_SB_; e.g., A55 in DENV2) is consistently shorter than the one connecting B and T stems (J_BT_; e.g., G17, G18, and A19 in DENV2). Finally, the 3WJ is strongly biased toward purine residues. Because both the sequence composition and the relative lengths of the junction strands govern the coaxial stacking of multi-way junctions (*46, 47*), the conservation of these features suggests that the conformational preferences of the 3WJ—and therefore the global structural dynamics of SLA—are conserved across flaviviruses.

To test this prediction, we used the scaffold-based cryo-EM approach described above to examine the 3D ensembles of SLAs from ZIKV, WNV, and YFV. DMS-MaPseq confirmed that the previously established secondary structures of TET and the SLAs are maintained in the scaffolded constructs (fig. S1). Despite differences in sequence, length, and 2-way junctions (fig. S10), the major 3D states of all SLAs are strikingly similar (Fig. 2C & 2D; fig. S11-13). A conformation consistent with the S-T state was well-resolved in all SLAs, producing 3D models where the 3WJs assume the same sharp bent configuration observed in DENV2 SLA (RMSD_backbone_ of 2.71 Å; Fig. 2E). Conformations consistent with the S-B state were also observed for all SLAs. They were well-resolved for ZIKV and YFV, but insufficiently well-defined for WNV to permit 3D structural modeling (fig. S12). Where modeled, the global structures and internal conformation of the 3WJ across S-B states are consistent (RMSD_backbone_ = 3.87 Å; Fig. 2F). For ZIKV, a minor fraction of the particles produced a 3D reconstruction that was consistent with a state in which the B and T stems stack to form a long semi-continuous helix (B-T state; fig. S11). A map consistent with the stacking of the B and T stems was also observed in YFV, but in this map the S stem was not visible suggesting that it is relatively more dynamic (fig. S13).

Overall, our cryo-EM comparisons strongly support the dynamic formation of S-T and S-B states across flaviviral SLAs. The observation of a possible B-T state suggests that the 3WJ may sample all three stacking configurations of the B, T and S stems but that the relative populations of these configurations differ across variants, in some cases being sufficiently populated to allow cryo-EM reconstruction. Indeed, all variants showed differences in particle distribution across states (fig. S14). For example, relative to the other SLAs, 3D classification analysis of WNV SLA assigned a larger fraction of the particles to unresolved 3D classes. This is consistent with the lower stability of the WNV SLA’s S stem—as previously observed (*48*) and confirmed by our DMS chemical probing experiments (fig. S1)—which increases the population of S stem-melted conformations. Thus, while the conformations accessible to the SLA are largely conserved, their relative populations are determined by sequence, S stem stability, and by solution conditions, such as [Mg^2+^].

Together, these results demonstrate that the 3D structural ensemble adopted by flaviviral SLAs is largely conserved and dictated by the intrinsic stacking propensities of the 3WJ. Multi-way junctions are known to adopt alternative stacking configurations (*49*), but how their dynamic interconversion encodes function is not well understood. The highly dynamic nature of the SLA, and its propensity to form S-B and S-T states across different flaviviral variants raised the question of whether these states and their interconversion play a role in the recognition and function of NS5.

### Conserved engagement of the S-T state by DENV and ZIKV NS5

We next asked how NS5 interacts with SLA 3D ensemble, and whether this is conserved across flaviviruses. To this end, we solved the cryo-EM structures of the DENV2 and ZIKV SLA-NS5 complexes that were reconstituted from recombinantly expressed and purified NS5, and *in vitro* transcribed 5′-capped SLA RNAs. Single-particle cryo-EM reconstructions yielded maps of 4.5 Å for DENV2 and 3.6 Å for ZIKV (Fig. 3A & 3B; fig. S15 & S16), after 3D refinement with Blush regularization (*50*). Comparison with the unbound ensembles revealed that the bound conformations of DENV2 and ZIKV SLAs closely match their S-T states (Fig. 3C & 3D), suggesting that NS5 specifically interacts with the S-T state of the unbound SLA conformational ensemble.

**Fig. 3.**
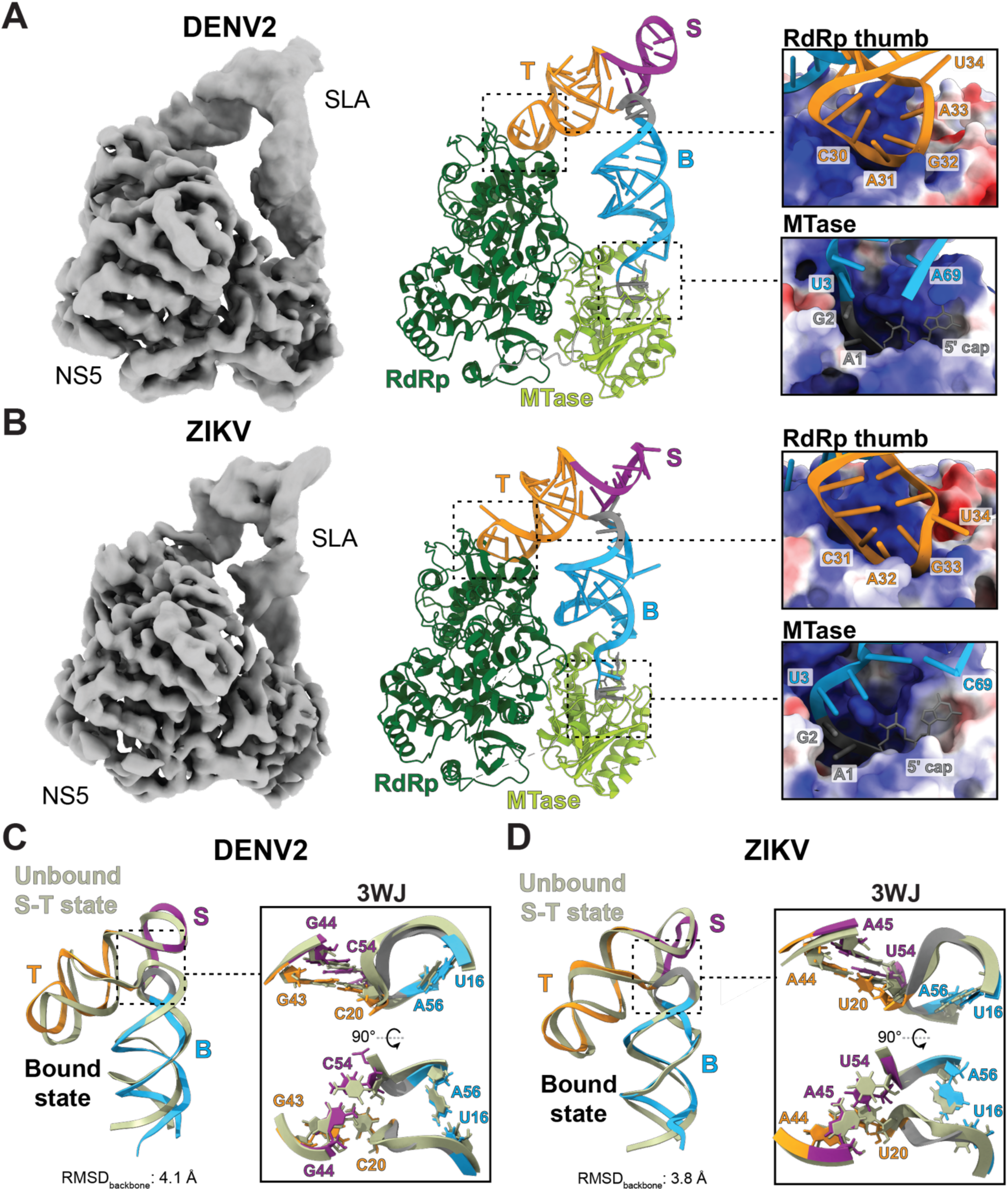
NS5 engages the S-T state of SLA in DENV and ZIKV. (**A**) Cryo-EM structure of DENV2 SLA-NS5 complex. The final reconstruction after 3D refinement is shown on the left, with the regions corresponding to SLA and NS5 labeled. Structural model of DENV2 SLA-NS5 complex (middle) with RdRp colored in dark green, MTase domain in light green, inter-domain linker in grey. SLA is colored using same color scheme as in Fig. 1A. SLA-NS5 complex forms two discrete RNA-protein interfaces (right). NS5 binding pockets are shown in surface representation colored by electrostatic potential, ranging from electronegative regions (red; -10 kcal/[mol·e]) to electropositive regions (blue; +10 kcal/[mol·e]), rendered using ChimeraX (*63*). SLA residues in the apical loop of the T stem (top right) and capped 5′ end (bottom right) contacting the RdRp thumb and MTase pockets, respectively, are highlighted. (**B**) Cryo-EM structure of ZIKV SLA-NS5 complex. The final map after 3D refinement (left), structural model of ZIKV SLA-NS5 complex (middle), and RNA-protein interfaces (right) are shown using same coloring and rendering scheme as in panel A. (**C**) NS5-bound DENV2 SLA adopts a conformation resembling the S-T state from its unbound ensemble. The bound (multicolor) and unbound S-T state (sage green) were superimposed via backbone atoms (left). Structural alignment of 3WJ using the closing base-pairs of the B, T and S stems labeled (right). (**D**) NS5-bound ZIKV SLA adopts a conformation resembling the S-T state from its unbound ensemble. The superposition of the bound and unbound S-T state (left) and structural alignment of 3WJ (right) are shown using the same color scheme as in panel C.

In contrast to previous proposals of species-specific SLA recognition modes (*35, 36*), the bound SLA conformations and the SLA-NS5 interfaces in the DENV2 and ZIKV complexes are nearly identical (Fig. 3A, B; fig. S17). In both complexes, the apical loop of the T stem makes contacts with NS5’s RdRp thumb subdomain and the 5′ end interacts with NS5’s MTase domain, resulting in two discrete RNA-protein interfaces. Moreover, both DENV2 and ZIKV complexes are nearly superimposable with a previously reported crosslinked DENV4 SLA-DENV3 NS5 complex (RMSD of 2.33 Å and 3.38 Å, respectively), in which the bound SLA likewise adopts the S-T conformation–consistent with the NS5-bound DENV2 and ZIKV SLA structures (RMSD_backbone_ of 3.61 Å and 3.55 Å, respectively) (fig. S17) (*51*). A structure of a crosslinked DENV2 SLA-NS5 complex has also been reported (*52*), but in that study the conformation of the bound SLA could not be determined directly from the density and a crystal structure of a dimerized DENV2 SLA was docked instead. Importantly, the 3WJ backbone configurations are nearly identical for the bound and unbound S-T states (Fig. 3C & 3D; fig. S17). Thus, three independent SLA-NS5 complexes converge on the same structure, strongly supporting a conserved mechanism of SLA recognition in which NS5 engages the S-T state of the 3D ensemble. This structural convergence is consistent with the functional interchangeability of SLAs among flaviviruses (*31–35*).

In addition to the density corresponding to the SLA-NS5 complex, our maps displayed a small region of undefined density near the RNA template entry channel of NS5 (fig. S18). This may be explained by interactions between a second SLA and this RNA-binding site on NS5, consistent with native mass spectrometry analysis indicating an SLA:NS5 stoichiometry of 2:1 at cryo-EM concentrations (fig. S19). This density was not observed in the crosslinked DENV4/DENV3 complex, likely due to crosslinking and additional purification steps in that study (*51*).

Together, these results establish that the S-T state of the SLA conformational ensemble represents a state that is specifically engaged by NS5, and that this complex is conserved across flaviviruses. Whether the conformational dynamics of SLA—and specifically the ability to populate the S-T state–are required for productive negative-strand synthesis remained to be investigated.

### In the S-T state, the 3WJ forms a ubiquitous 3D motif

Having established that NS5 engages the S-T state, we sought to understand the molecular interactions within the 3WJ that stabilize this conformation. Our DENV2 and ZIKV SLA-NS5 complexes and the unbound S-T structures provided the global architecture of the 3WJ, but the moderate resolution of these structures limited our ability to resolve the stabilizing interactions at atomic detail. Because the 3WJ conformations of our bound and unbound structures are nearly superimposable with that of the bound DENV4 SLA in the crosslinked complex (fig. S17), we leveraged the higher resolution of that structure to examine atomic details within the 3WJ (Fig. 4A).

**Fig. 4.**
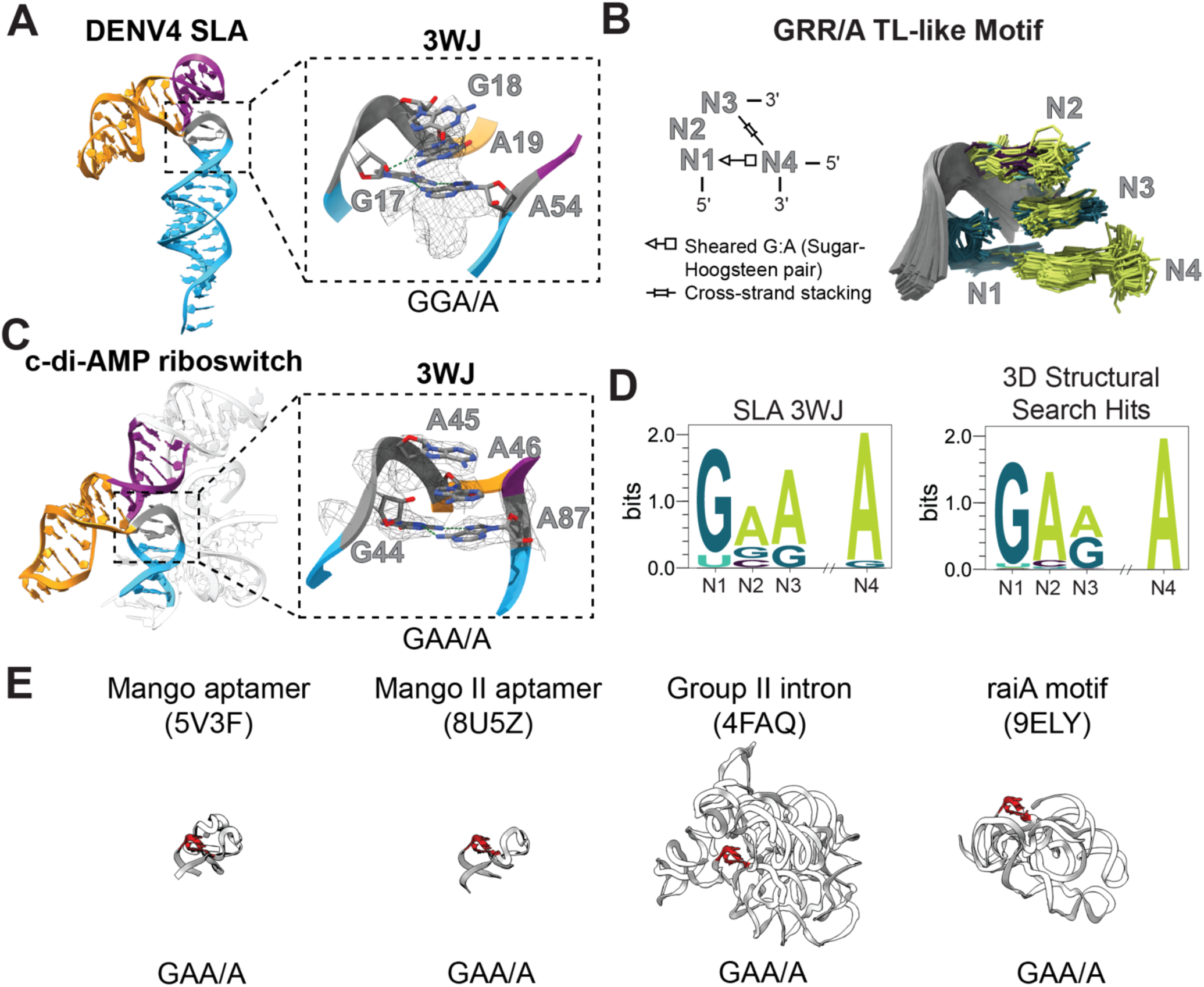
SLA 3WJ in S-T state forms a ubiquitous 3D motif. **A**) 3WJ configuration of NS5-bound DENV4 SLA (PDB: 8GZP) (*51*). The 3WJ residues are labeled according to DENV4 numbering, with the cryo-EM density overlaid. **B**) GRR/A TL-like motif. The motif topology, including the cross-strand stacking interaction and sheared G:A base pair, is illustrated in the left diagram. The structural alignment of all the hits from our PDB search is shown on the right. Bases are colored according to logo plots in panel D. **C**) c-di-AMP riboswitch helical arrangement around the 3WJ with the GRR/A TL-like motif (PDB: 4QLM) (*64*). 3WJ motif with the corresponding density is highlighted on the right. **D)** Sequence conservation of the SLA 3WJ generated from the sequence alignment in Fig. 2A (left). Sequence logo of GRR/A TL-like junction motif generated from all WebFR3D hits (right). **E**) Examples of WebFR3D hits of GRR/A TL-like motif in junctions. GRR/A TL-like motif is shown in red with the corresponding motif sequence at the bottom.

Inspection of the NS5-bound DENV4 SLA revealed that the conformation of the 3WJ is stabilized by a sheared G:A base pair formed by G17 and A54, a cross-strand stacking between A54 and A19, and the stacking of G18 with A19 (Fig. 4A). These interactions produce the sharp ∼180° bend of the RNA backbone required for the coaxial stacking of the S and T stems. Notably, sheared G:A base pairs are hallmarks of sharp backbone kinks that are critical for the folding of complex 3D structures (*53–55*). The specific arrangement observed in the 3WJ—a sheared G:A pair coupled with cross-strand purine stacking—has been previously observed in the junctions of Mango aptamers and the raiA bacterial non-coding RNA, where it was noted to resemble an interrupted GNRA TL, assembled from non-contiguous residues (*56, 57*). A systematic search of the Protein Data Bank (PDB) using the WebFR3D server revealed that this configuration is widely distributed across functional RNAs, including group II introns, aptamers, ribosomal RNAs, and viral tRNA-like structures (Fig. 4B; Supplementary Data 1). Many of the examples found in our search represent tertiary interactions where distant adenines form sheared base pairs with guanines within T-loops (fig. S20). In many other examples—such as in the SLA S-T state—the motif is found inside of junctions, where it directs the relative orientation of the emanating helices, as is the case for the c-di-AMP riboswitch where the helical arrangement mirrors that of the SLA (Fig. 4C). Recently, a sequence alignment-based probabilistic RNA grammar suggested that this motif is highly ubiquitous at three-way junctions (*58*). Thus, the 3WJ conformation observed in the S-T state corresponds to a ubiquitous 3D motif that can form in distinct structural contexts and that stabilizes the 3D structure of diverse RNAs.

A sequence logo generated from the variants identified from our 3D structural search mirrors that of SLA’s 3WJ, indicating a strong preference for a sheared G:A base pair (at positions N1 and N4) with two purine (R) residues (primarily adenosines) at positions N2 and N3 (Fig. 4D). Based on these sequence preferences, we refer to it as the GRR/A TL-like motif hereafter. This strongly suggests that the sequence of SLA’s 3WJ is evolutionarily constrained to fold into the GRR/A TL-like motif, presumably to stabilize the S-T state required for engagement with NS5 and negative-strand synthesis. In other RNAs where the motif was identified inside junctions (Fig. 4E), comparisons of aligned variants displayed similar sequence preferences for the motif (fig. S20).

Although RNA 3D motifs are typically considered as rigid building blocks of RNA 3D architecture, our cryo-EM and smFRET data indicate that the SLA’s 3WJ dynamically samples a conformation where the GRR/A TL-like motif is formed (the S-T state) and conformations in which it is not (the S-B state and others). Because this conformational sampling shapes the global 3D ensemble of the SLA, we next investigated whether and how it impacts NS5 recognition and function.

### 3WJ ensemble guides assembly of a replication initiation-competent complex via a post-binding conformational search

Because the sheared G:A base pair is conserved across the 3WJs of SLA and is central to the GRR/A TL-like motif that stabilizes the S-T state, we investigated how perturbing this interaction affects the SLA ensemble and its ability to bind NS5 and promote negative-strand synthesis. In DENV2 SLA, this base pair is formed between G17 and A55 (Fig. 1A). We designed a DENV2 SLA construct containing a single G17A mutation that disrupts the G:A base pair and characterized its effects on the SLA ensemble by smFRET (Fig. 5A). After confirming that the mutation does not affect the overall folding of SLA (fig. S21), smFRET revealed that, unlike wild-type (WT)—in which both the S-T (high FRET) and S-B (low FRET) states are clearly resolved—the G17A mutant predominantly populates the S-B state across solution conditions (Fig. 5A; fig. S21). Thus, the G17A mutation strongly destabilizes the S-T state, shifting the global 3D ensemble towards the S-B state. Consistent with this, our motif search returned no AGA/A hits matching the G17A sequence (Supplementary Data 1), further supporting that the mutation disfavors formation of the GRR/A TL-like motif.

**Fig. 5.**
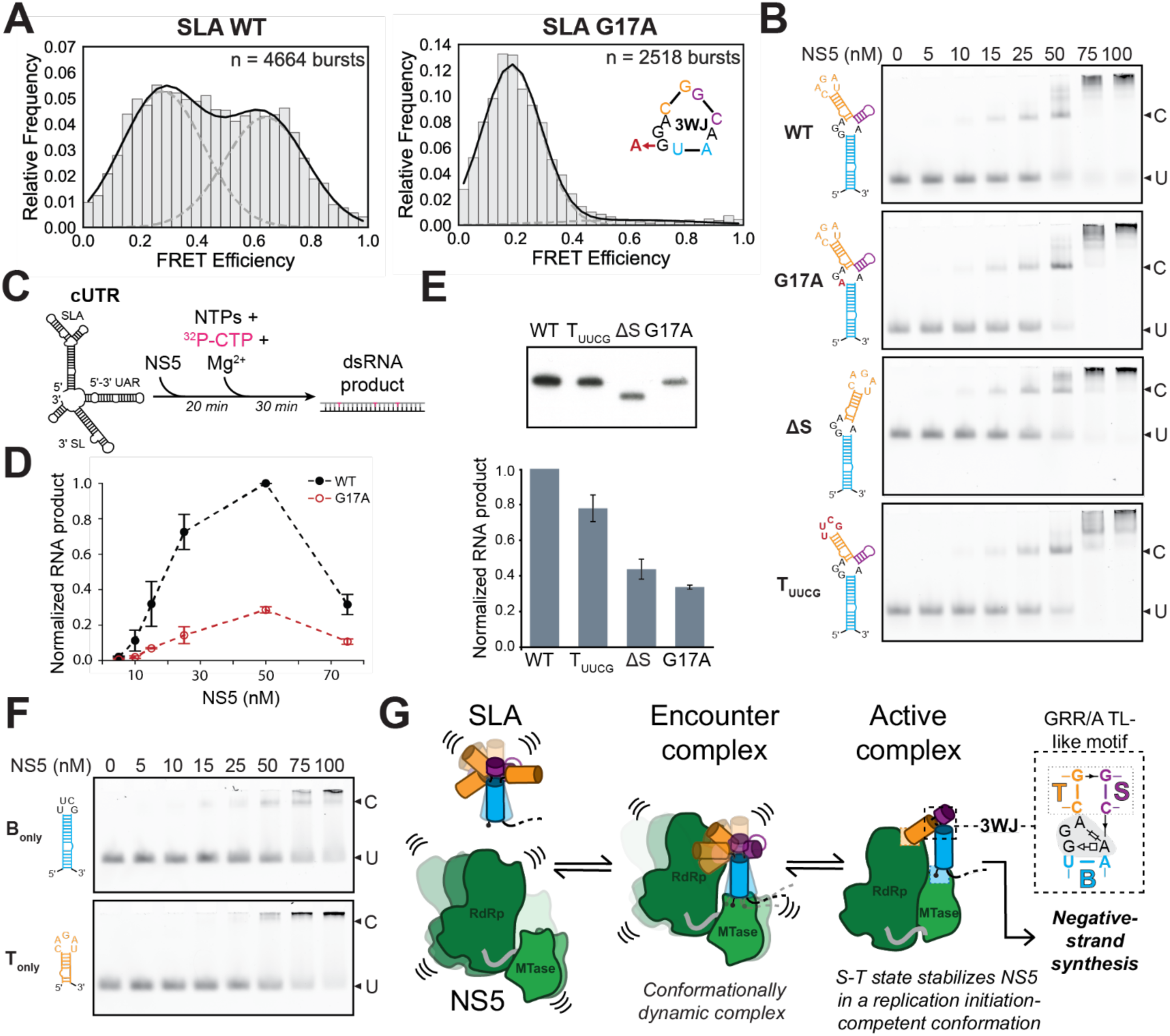
SLA 3D ensemble modulates NS5 function via a post-binding conformational search. **A**) Normalized FRET efficiency histogram at 5 mM MgCl_2_ is shown for WT and G17A SLA, fitted to a two-Gaussian model. **B**) NS5 binding to 3′ Cy3-labeled SLA variants by EMSA. Bands corresponding to unbound SLA (U) and SLA-NS5 complex (C) are labeled. **C**) Experimental scheme for radioactivity-based negative-strand synthesis from a truncated, circularized RNA template (cUTR). cUTR and NS5 are incubated for 20 min before the reaction is initiated by adding NTPs, 32P-CTP and Mg^2+^. Reaction is quenched after 30 min. **D**) Titration of NS5 for negative-strand synthesis. Quantification of ^32^P-CTP labeled full-length product by densitometry analysis of fig. S22. **E)** Negative-strand synthesis activity for different SLA variants. Full-length RNA products (top) and their quantification by densitometry analysis (bottom) are shown. **F**) NS5 binding to B stem (B_only_) and T stem (T_only_) SLA fragments evaluated by EMSA. The bands corresponding to unbound RNA (U) and RNA-NS5 complex (C) are indicated. **G**) Proposed model of post-binding conformational search of SLA-NS5 complex leading to negative-strand synthesis. The flexibility of NS5 provided by its 10-amino acid linker allows it to accommodate multiple SLA conformations in an initially dynamic encounter complex. A conformational search in which SLA adopts the S-T state results in the stabilization of the active SLA-NS5 conformation that is competent for initiation of negative-strand synthesis.

Next, we measured how perturbing the DENV2 SLA 3D ensemble—specifically its ability to sample the S-T state—affects NS5 binding. In addition to WT and G17A SLA, we included a mutant in which the S stem was deleted (ΔS). This mutation converts the 3WJ into a two-way junction but retains the residues involved in formation of the GRR/A TL-like motif, which may still form locally, albeit with reduced stability from the loss of S stem stacking. We also generated another mutant where the CAGAU apical loop of the T stem was replaced with a UUCG tetraloop (T_UUCG_). In the SLA-NS5 cryo-EM structure, this apical loop makes contacts with NS5’s RdRp thumb subdomain (Fig. 3A). Thus, we predicted that the T_UUCG_ mutation would perturb a direct RNA-protein contact while not affecting the SLA’s 3D ensemble. We assessed SLA-NS5 binding by electrophoretic mobility shift assays (EMSA).

For WT SLA, we observed NS5 concentration-dependent SLA-NS5 complex formation, with the majority of SLA bound at 50 nM NS5 (Fig. 5B). As reported in previous studies (*31, 35*), higher-order complexes were also observed; they became predominant at [NS5] > 50 nM (Fig. 5B). Given that the S-T state is observed in the SLA-NS5 cryo-EM structures, we expected that its destabilization would lead to weaker NS5 binding, consistent with a conformational capture model. Surprisingly, in contrast with this model, NS5 binding of the G17A and ΔS mutants was essentially identical to that of WT SLA (Fig. 5B), despite the perturbations to the 3WJ that substantially destabilize the S-T state. The T_UUCG_ mutant also showed comparable binding to WT (Fig. 5B), suggesting that contacts between the apical loop and the RdRp thumb subdomain either are not sequence-specific or form after initial SLA-NS5 binding. The absence of a binding effect upon destabilization of the S-T state or mutation to T_UUCG_ suggests that the SLA-NS5 conformation observed by cryo-EM corresponds to a state that forms after initial binding—i.e., multi-step recognition—or alternatively, that the observed cryo-EM structure does not represent a functional state. To distinguish between these possibilities, we tested the effect of these mutations on negative-strand synthesis.

We synthesized minimal constructs that mimic the circularized form of the DENV2 genome (cUTR), containing the SLA, the 5′-3′ upstream AUG region (UAR) complementary sequences, and 3′ stem-loop (SL) (Fig. 5C). Similar constructs have been used previously for *in vitro* negative-strand synthesis assays and structural studies (*15, 51*). We synthesized cUTR variants harboring WT, G17A, ΔS and T_UUCG_ SLAs and assessed their ability to bind NS5 by EMSA. Consistent with our previous results, NS5 binding was comparable across all cUTR variants (WT, G17A, ΔS and T_UUCG_)—i.e., the mutations to SLA did not have a measurable effect on NS5 binding (fig. S22). The NS5-binding profiles of the cUTR constructs closely resembled those of the isolated SLAs, with most RNA bound at 50 nM NS5 (Fig. 5B; fig. S22). This indicates that interactions between NS5 and cUTR are predominantly driven by SLA-NS5 binding.

We assessed negative-strand synthesis activity by quantifying full-length negative-strand product formation using radiolabeled NTP incorporation (Fig. 5C). Negative-strand synthesis increased with increasing amounts of NS5—proportional to the formation of the SLA-NS5 complex observed in our binding assays—confirming that the SLA-NS5 interactions we observed reflected functional interactions and that the cUTR templates are competent for SLA-dependent negative-strand synthesis (Fig. 5D; fig. S22). At NS5 concentrations where higher-order complexes or aggregates predominate (> 50 nM), we observed a reduction in negative-strand synthesis, consistent with the loss of the functional SLA-NS5 complex.

Despite showing no effect on NS5 binding, the mutations substantially attenuated NS5-catalyzed negative-strand synthesis. Notably, the single-point mutant G17A that destabilized the S-T state (Fig. 5A) significantly reduced negative-strand synthesis across all NS5 concentrations tested (Fig. 5D). At 50 nM NS5, where we observed maximum negative-strand synthesis activity, the G17A mutation reduced negative-strand synthesis by 66%, while the ΔS mutation reduced it by 56% (Fig. 5E). The T_UUCG_ mutant reduced negative-strand synthesis by 22% (Fig. 5E). Thus, disruption of the 3WJ’s ensemble and its ability to form S-T state, can have a large impact on negative-strand synthesis and these effects were larger than the perturbation to the RNA-protein interface produced by the T_UUCG_ mutant. This underscores the functional importance of the 3WJ and SLA’s conformational ensemble.

The contrasting effects of the SLA’s 3WJ mutations—no effect on NS5 binding but a large effect on negative-strand synthesis—suggest that the S-T state forms after an initial “encounter complex” in which SLA binds NS5 without a strict requirement for a specific SLA 3D conformation. This could reflect either the formation of only one of the two RNA-protein interfaces identified in the cryo-EM complex, or the participation of both interfaces in a manner that accommodates multiple SLA 3D conformations through the combined flexibility of NS5 and SLA.

To distinguish between the above possibilities, we designed truncated SLA constructs predicted to contain only one of the two NS5-binding interfaces: a T_only_ construct containing the T stem and apical loop, predicted to engage NS5’s RdRp thumb subdomain, and a B_only_ construct, predicted to engage the MTase domain (Fig. 5F). Both T_only_ and B_only_ bound NS5 substantially more weakly than full SLA (Fig. 5F; fig. S22). In contrast, the ΔS construct, which contains both interfaces, bound similarly to WT SLA. Together, these observations indicate that NS5 engages both the T and B stems of SLA in the encounter complex. Relative to their positions in the cryo-EM complex, however, these contacts may be only partially formed, be flexible, and/or adopt variable geometries. Consistent with this, the RdRp and MTase domains of NS5 are connected by a flexible ∼10-residue linker that permits substantial interdomain motion in solution (*59–62*), allowing both interfaces to be satisfied across a range of relative domain orientations. Additionally, contacts between SLA and the MTase domain are mediated by single-stranded 5’ end of the SLA (Fig. 3A), which is intrinsically flexible and may therefore tolerate variation in the global 3D conformation of the SLA.

Together with the mutational analysis described above, our results support a multi-step recognition mechanism involving conformational changes of the RNA after initial NS5 binding (Fig. 5G). Although previous studies observed mutations that affect negative-strand synthesis without affecting NS5 binding, suggesting a post-binding step (*15, 33*), the underlying mechanism remained mysterious. Our results provide a structural and mechanistic framework for this step. In the encounter complex, the conformational flexibility of both NS5 and SLA enables engagement across multiple SLA conformations—explaining why perturbations to the global 3D ensemble do not affect binding. After initial association, a subsequent conformational search, mediated by motions in both the RNA and the protein, drives formation of the S-T state captured in our cryo-EM structures. Stabilized by the GRR/A TL-like motif and coaxial stacking interactions, the S-T state positions the MTase and RdRp domains of NS5 in a conformation that can engage the 3’ initiation site and initiate negative-strand synthesis. Thus, the intrinsic conformational dynamics of the SLA mediate the progression from a flexible encounter complex to a state competent for replication initiation.

## DISCUSSION

A central challenge in RNA biology is to understand how conformational dynamics contribute to RNA function. Although RNA structures have long been recognized as regulators of critical viral processes and as potential therapeutic targets, structural studies of these elements have largely treated them as static, limiting our mechanistic understanding of how they are recognized by and function with their protein partners. The SLA promoter in flaviviruses exemplifies this gap: previous structural studies captured divergent conformations depending on the specific SLA variant and technique used, and implicitly treated it as a passive binding site for NS5 (*28, 35*), despite evidence for local structural dynamics (*29*). The divergent structures, rather than reflecting virus-specific recognition modes, instead represent distinct states within a conserved, dynamic conformational ensemble. This conservation in the SLA 3D structural ensemble supports a shared mechanism of NS5 recognition, consistent with the functional interchangeability of SLAs across flaviviruses (*31–35*).

By adopting an ensemble-based perspective—integrating scaffold-aided cryo-EM, smFRET, and biochemical assays—we show that SLAs from DENV2, ZIKV, YFV, and WNV populate a conserved set of globally distinct 3D conformations that are governed by alternative stacking configurations of the B, T, and S stems. These configurations are in turn dictated by the intrinsic structural dynamics of the central 3WJ, which has shared properties across variants—including rich purine composition and strand length asymmetries. Our cryo-EM structures of DENV2 and ZIKV SLA-NS5 complexes, along with a previously reported crosslinked DENV4/DENV3 SLA-NS5 structure, demonstrate that NS5 recognizes a specific state of the unbound SLA ensemble: the S-T state, in which S and T stems coaxially stack. Together, three independent SLA-NS5 structures converge on a conserved recognition mode.

Central to the S-T state is the GRR/A TL-like motif—a configuration stabilized by a sheared G:A base pair and a network of stacking interactions. A systematic search of the PDB reveals that this motif is broadly distributed across diverse functional RNAs, in many cases as part of T-loop mediated tertiary interactions and in other cases as part of junctions—as with the SLA’s S-T state—directing the relative orientation of emanating helices. Although RNA 3D motifs are often considered rigid building blocks, the GRR/A TL-like motif is conformationally dynamic, and in the SLA this flexibility guides a post-binding conformational search for the replication initiation-competent SLA-NS5 complex. Thus, understanding the intrinsic conformational dynamics of recurring RNA 3D motifs is critical for dissecting RNA structure-based mechanisms and for building predictive models of RNA structural dynamics and function.

In DENV2 SLA, a mutation that disrupts the conserved G:A base pair of the GRR/A TL-like motif destabilizes the S-T state and greatly reduces negative-strand synthesis. Despite these effects on the SLA conformational ensemble and on NS5 function, the mutation does not have a measurable effect on NS5 binding. Although a decoupling between SLA-NS5 binding and RdRp activity has been previously noted based on biochemical and functional studies (*15, 33*), the mechanistic basis for this observation remained unclear. Our results support the following mechanism: the lack of a binding effect of disrupting the GRR/A TL-like motif—despite its substantial effect on the SLA conformational ensemble—rules out a simple, one-step conformational-capture mechanism. Instead, they support a multi-step recognition pathway in which NS5 and SLA first associate into a conformationally flexible encounter complex and then undergo a post-binding conformational search that drives SLA into the S-T state and the complex into a replication initiation-competent conformation. Completing negative-strand synthesis initiation requires that NS5 engages the 3’ initiation site—a step that likely involves interactions with 3’ UTR structures and additional conformational changes in the RNA template and NS5 itself (*13, 14*). How these subsequent steps are coordinated remains an important open question that the mechanistic framework established here now makes tractable.

Beyond the mechanistic insights, our findings raise the possibility that the SLA’s conformational dynamics could inform future strategies to disrupt flavivirus replication. In principle, ligands that destabilize the S-T state–for example, by perturbing the GRR/A TL-like motif within the 3WJ–could interfere with the post-binding conformational search required for productive replication initiation, without requiring direct competition for NS5 binding. Consistent with the SLA being amenable to chemical perturbation, small molecules that bind the SLA have recently been reported (*33*). Whether the SLA’s intrinsic dynamics can be modulated in cellular contexts to disrupt replication remains to be tested, but the structural and mechanistic framework established here defines the relevant conformational landscape and identifies the structural dynamics of the 3WJ as a potential target for further investigation.

## Acknowledgments

We thank M. Ebrahim, J. Sotiris, and H. Ng for their assistance with cryo-EM experiments at the Evelyn Gruss Lipper Cryo-Electron Microscopy Resource Center. We thank J. Banfelder, L. Sweezy, B. Jayaraman, and R. Bennett for their assistance at the High-Performance Computing Resource Center. We thank R. Roeder for his generosity with equipment necessary for negative-strand synthesis assays. We thank J. Mendez with his assistance with cryo-EM experiments at NYSBC. We thank M. Abdelhamid for his feedback on smFRET experiments and data analysis. We thank C. Langeberg, J. Kieft, and members of the Bonilla lab for reading and providing feedback on the manuscript.

## Author contributions

L.V.S. conceptualized research, performed cryo-EM, smFRET and biochemical experiments, analyzed data, structural modeling, performed bioinformatic analysis, and wrote manuscript in collaboration with S.L.B. T.O. performed biochemical experiments, purified proteins, analyzed data, assisted with structural studies, and edited manuscript. M.A.A. performed 3D motif search analysis, performed bioinformatic analysis, and edited manuscript. E.A.C. assisted with RNA synthesis and molecular biology experiments. P.D.B.O. performed native mass-spec experiments and edited manuscript. L.U. assisted with cryo-EM experiments and structural modeling, and edited manuscript; D.I. performed DMS-MaPseq experiments and edited manuscript. S.L.B. conceptualized research, supervised all experiments and data analyses, assisted with experiments, wrote manuscript in collaboration with L.S.V., and obtained funding.

## Data, code, and materials availability

Cryo-EM maps and structural models will be deposited in the Electron Microscopy Data Bank (EMDB) and Protein Data Ban (PDB) upon publication. Motion corrected micrographs and particles will be deposited in Electron Microscopy Public Image Archive (EMPIAR) and will become available upon publication.

