## Supplementary Information for "RNA 3D Motif Dynamics Guide Assembly of the Replication Initiation Complex in Flaviviruses"

Lorena V. Streit *et al.*

**The PDF file includes:**

Materials and Methods

Figs. S1 to S22

Tables S1 to S4

**Other Supplementary Material for this manuscript includes the following:**

Supplementary Data 1

### MATERIALS AND METHODS

#### *In vitro* transcription, purification, and capping of RNA constructs

Templates for *in vitro* transcription were purchased as double-stranded DNA gene fragments (gBlocks; Integrated DNA Technologies) and amplified by PCR. The sequences for all constructs are provided in Table S2. *In vitro* transcription was performed using T7 RNA polymerase in reactions containing 6 mM NTPs (ATP, UTP, GTP, and CTP), 60 mM $MgCl_2$, 30 mM Tris-HCl pH 8.0, 10 mM DTT, 2 mM spermidine, 0.01% (v/v) Triton X-100, 6 U RNasin Plus RNase inhibitor (Promega), and ~0.14 mg/mL T7 RNA polymerase (prepared in-house). The transcription reactions were incubated at 37°C for 3 h. To produce homogeneous 3' ends in cUTR constructs used for negative-strand synthesis assays, a self-excising HDV ribozyme was appended immediately downstream of the cUTR sequence, as done previously (*65-67*). In those cases, the 3 h transcription reaction was followed by increasing [$MgCl_2$] to 125 mM and incubating at 60°C for 15 min. The transcribed RNA constructs were ethanol precipitated, purified by denaturing gel electrophoresis, and resuspended into RNase-free water. The purity of the RNAs was confirmed by denaturing electrophoresis. RNA folding was assessed by native polyacrylamide gel electrophoresis. To remove the 2′,3′-cyclo-phosphate group resulting from the self-excision of the HDV ribozyme from the cUTR constructs, 1 μM of RNA was incubated with 60 U of T4 polynucleotide kinase (M0201S, New England Biolabs) in 500 μL (100 mM Tris pH 6.5, 100 mM Mg acetate, 5 mM β-mercaptoethanol) at 37°C for 6 h. The RNA was purified by phenol-chloroform extraction and buffer exchanged into water.

For structural studies of the SLA-NS5 complexes, DENV2 and ZIKV SLA were 5' capped with a type-1 cap. DENV2 SLA was capped co-transcriptionally using a trinucleotide cap analog (E2080S, New England Biolabs) and ZIKV SLA was capped post-transcriptionally after purification via one-pot reaction with Faustovirus capping enzyme and Cap 2′-O-methyltransferase, as per manufacturer instructions (M2081 & M0366, New England Biolabs). The enzymatically capped SLA was phenol-chloroform extracted, ethanol precipitated, and buffer exchanged into RNase-free water. Capping of both SLAs was confirmed by denaturing polyacrylamide gel electrophoresis and native mass spectrometry.

**DMS chemical probing of RNA secondary structure**

10 µg RNA diluted in 20 µL nuclease-free water was supplemented with 5 µL RNA Folding Buffer 5X (250 mM Tris-HCl pH 7.5, 50 mM $MgCl_2$), heated at 50°C for 30 min and then allowed to cool to room temperature for 10 min. Dimethyl sulfate (DMS) (D186309, Merck) was diluted 1:6 in ethanol, added to the RNA at a final concentration of 100 mM, and allowed to react for 5 min at room temperature. Reactions were quenched by addition of 1 volume 1 M DTT, followed by RNA purification on Monarch® Spin RNA Cleanup columns (10 µg) (T2030L, New England Biolabs).

**DMS-MaPseq library preparation**

Library preparation was conducted as previously described (*68*), with minor changes. Briefly, 500 ng probed RNA were combined with 0.5 µL of 100 µM random hexamers, 0.5 µL of deoxynucleoside triphosphates (dNTPs; 10 mM each) and 1 µL of 5× RT buffer (250 mM Tris-HCl pH 8.3, 375 mM KCl and 15 mM $MgCl_2$). Samples were then incubated at 94°C for 5 min to simultaneously denature and fragment the RNA and immediately transferred to ice for 1 min. Samples were then supplemented with 0.25 µL of 0.1 M DTT, 5 U of SUPERase•In RNase inhibitor (A2696, Thermo Fisher Scientific), 50 U of Induro® Reverse Transcriptase (M0681L, New England Biolabs) and incubated at 25 °C for 10 min, 55 °C for 30 min and 60 °C for 30 min, followed by heat inactivation at 75°C for 15 min. cDNA was then converted to double-stranded DNA (dsDNA) using the NEBNext Ultra II Non-Directional RNA Second Strand Synthesis Module (E6111L, New England Biolabs) by incubating at 16 °C for 1.5 h. dsDNA was cleaned up with 1.8 volumes of NucleoMag NGS cleanup and size select beads (744970, Macherey Nagel) and used as input for the NEBNext Ultra II DNA library prep kit for Illumina (E7645S, New England Biolabs) as per manufacturer instructions, but scaling down all reaction volumes 4-fold.

**DMS-MaPseq data analysis**

Reads were mapped the *rf-map* tool of RNA Framework (*69*) version 2.8.6 and Bowtie2 (*70*) version 2.3.5.1 after clipping terminal bases with Phred quality < 20, discarding reads containing internal Ns and trimming terminal Ns (parameters: *-b2 -cq5 20 -ctn -cmn 0 -mp '--very-sensitive-local'*). Alignments in SAM format were sorted and converted to BAM format using SAMtools (*71*) version 1.15.1. BAM alignments were then processed using RNA Framework's *rf-count* to generate RC files (containing per-base mutations and coverage) (parameters: *-m -na -ni -ds 90 -ncl -dc 3 -me 0.1*). RC files were then processed using RNA Framework's *rf-norm* to obtain normalized reactivity profiles (parameters: *-sm 4 -nm 3 -rb AC -mm 1 -n 1000*).

**Plasmid construction**

DENV2 and ZIKV NS5 gene fragments were codon-optimized for *E. coli* expression and ordered as gBlocks (Integrated DNA Technologies; sequences are shown in Table S2). The gene fragments were amplified by PCR and cloned into the pET15b vector downstream of the thrombin digestion site using the NEBuilder HiFi DNA Assembly Cloning Kit (E2621, New England Biolabs). The resulting constructs contained an N-terminal 6x His-tag followed by a thrombin digestion site, linker and NS5.

**NS5 expression and purification**

DENV2 and ZIKV NS5 were both expressed in BL21(DE3) cells, and the same protocol was used for purification. BL21(DE3) cells transformed with pET15b-NS5 were grown to $OD_{600}$ of 0.4 at 37°C. Then, the culture was grown for another hour at 18°C before inducing NS5 expression with 0.4 mM IPTG and was grown overnight before harvest.

The cells were lysed in NS Buffer (20 mM HEPES pH 8.0, 10% (v/v) glycerol, 5 mM DTT) with 400 mM NaCl by sonication and cleared by centrifugation. The lysate was loaded onto Ni-NTA beads (Qiagen) in an open column (Bio-rad) by gravity flow, and the beads were successively washed with NS Buffer with 400 mM NaCl containing 0, 25, 50, and 75 mM imidazole, and NS5 was eluted with NS Buffer containing 400 mM imidazole. The eluate was dialyzed into NS Buffer to remove imidazole, and NS5 was treated with thrombin (T4648, Sigma) and subsequently purified over HiTrap Heparin in NS Buffer with a salt gradient from 400 mM and 200 mM NaCl for DENV2 and ZIKV, respectively, to 900 mM NaCl. Finally, NS5-containing fractions were further purified using Superdex 200 16/60 (Cytiva) pre-equilibrated in NS Buffer with 400 mM NaCl. Fractions containing NS5 were collected and concentrated using Amicon concentrator with 10 kDa MWCO (UFC901024, Millipore). The protein was snap frozen and stored at -80°C.

**Electrophoretic mobility shift assays (EMSAs)**

Two different types of constructs were evaluated for binding to NS5; ~70-nt Cy3-labeled SLA and ~180-nt cUTR. 20 nM SLA-Cy3 or cUTR constructs were refolded by heating at 80°C for 1 min or 50°C for 5 min, respectively, and cooling to room temperature for 10 min in 10 mM HEPES pH 8.0. The refolded RNA was mixed with 10× NS5 (20 mM HEPES pH 8.0, 150 mM NaCl, 5 mM DTT, 10% (v/v) glycerol) in 20 mM HEPES pH 8.0, 10 mM KCl, 10 mM DTT and 50 µg/mL BSA (AM2616, Invitrogen) at a final RNA concentration of 10 nM in 10 µL. After the reaction was incubated for 30 min at 30°C, 2 µL of 24% (v/v) glycerol was added for sample loading and the mixture was resolved on 4.5% (w/v) polyacrylamide-TBE gels with 4% (v/v) glycerol. For the cUTR constructs, the gels were stained with SYBR Gold (S11494, Invitrogen) and images were taken using ChemiDoc (Bio-Rad) with auto-optimal setting. For SLA-Cy3, the gel image was taken directly after gel running with the Cy3 channel of ChemiDoc without any staining. For quantification, densitometric analyses were performed on scanned gel images using Imagequant TL (Cytiva). The band intensities of free RNA were determined for each concentration, normalized to 0 nM NS5, and means and standard deviations were calculated from three replicates.

***In vitro* negative-strand synthesis assays**

*In vitro* negative-strand synthesis assays were set up as for EMSA, as described above, in the final volume of 25 µL. After mixing cUTR templates with NS5, the reaction mixture was incubated for 20 min at 30°C. Then, NTPs were added to the final concentrations of 0.5 mM each of ATP, GTP, UTP, and 0.025 mM CTP together with 5 µCi of $^{32}$P-α-CTP (BLU008H, Revvity). RNA synthesis was then initiated by adding $MgCl_2$ solution to 5 mM, and the reaction was carried out for 30 min at 30°C. The reaction was terminated with Stop Buffer (1.28% (w/v) SDS, 23.1 mM HEPES pH 8.0, 23.1 mM EDTA, 231 mM NaCl, and 0.429 mg/ml yeast tRNA), and RNA was purified by ethanol precipitation following phenol chloroform treatment. Precipitated RNA was resuspended in RNA sample buffer containing 80% formamide and boiled for 5 min at 95°C. RNA was resolved on a 6% (w/v) acrylamide-TBE gel with 8 M urea, and RNA was detected using film autoradiography. For quantification of RNA products, densitometric analyses were performed on scanned images of films using Imagequant TL (Cytiva). Multiple time points were used to expose films, and the time point that produced a densitometer read of about 1,500,000 for 50 nM NS5 and WT cUTR, which showed the strongest activity in our assays, was used (either 45 min or 1 h exposure times). Densitometer reads of all RNA products were normalized to the read of WT cUTR at 50 nM NS5 from the same film, and means and standard deviations were calculated from three experiments. An exposure time producing a densitometric read of approximately 1,500,000 for the 50 nM NS5 WT condition was selected as it represented the maximal exposure without significant signal saturation.

**Cryo-EM sample preparation, grid freezing, and data collection**

DENV2 SLA and all scaffolded constructs were refolded in 50 mM Tris-HCl pH 7.5 and 10 mM $MgCl_2$ by heating at 50°C for 30 min and cooling to room temperature for 10 min. C-Flat holey carbon grids (hole size: 1.2 µm; spacing: 1.3 µm; mesh: 400; Electron Microscopy Sciences) were cleaned with a Gatan Solarus 950 advanced plasma system using atmospheric air plasma (70 mTorr) for 15 sec. 2.5 µL of the refolded RNA sample (75 µM of DENV2 SLA and 20 µM of SLA-TET/cpTET) were deposited on the grid at 4°C and 100% humidity and subsequently plunge-frozen in liquid ethane in a FEI Vitrobot 3 (blot time: 1-2 sec; blot force: 0; wait time: 0).

For the DENV2 and ZIKV SLA-NS5 complexes, the 5' capped SLAs were refolded in 50 mM Tris-HCl pH 7.5 and 5 mM MgCl2 by heating at 80°C for 1 min and cooled to room temperature for 10 min. The complexes were assembled at room temperature for 30 min in 50 mM Tris-HCl pH 7.5, 75 mM NaCl, 1 mM MgCl2, 5 mM DTT, with glycerol at 2.5% (v/v) and 5% (v/v) for ZIKV and DENV2 complexes, respectively.

2.5 µL of 2.5 µM DENV2 SLA-NS5 complex was deposited on C-Flat holey carbon grids (hole size: 1.2 µm; spacing: 1.3 µm; mesh: 400; Electron Microscopy Sciences) and 2.5 µL of 2 µM ZIKV SLA-NS5 complex was deposited on QuantiFoil holey carbon grids (hole size: 2 µm; spacing: 1 µm, mesh: 200; Electron Microscopy Sciences). The grids were glow-discharged with a Gatan Solarus 950 advanced plasma system using atmospheric air plasma (70 mTorr) for 15 sec.

Grid screening for all datasets was performed in a Thermo Fisher Scientific Talos Arctica transmission electron microscope (TEM) operated at 200 kV and equipped with either a Gatan K2 or a Gatan Alpine direct electron detector camera. All datasets were collected on a Thermo Fisher

Scientific Krios TEM operated at 300 kV at a defocus range of -0.7 to -2.5 µm, with a total dose of 50-60 e/Å$^2$. Sample and data collection parameters are summarized in Table S4.

**Cryo-EM data analysis**

Cryo-EM data were processed using cryoSPARC (v. 4.5.3) (*72*) and RELION (v. 5.0.0) (*73*). The processing pipeline for each dataset is summarized in Figures S2, S4, S6, S11-13, S15, S17. In brief, movies were motion corrected using 2× binning for all datasets (patch or global motion correction, as specified per dataset) and CTF estimation was performed using default parameters. Micrographs with obvious defects were excluded using exposure curation. Particles were picked using different methods depending on the dataset, including blob, template and Topaz picker v. 0.3.10 (*74*). Extracted particles were cleaned using multiple rounds of 2D classification to generate initial *ab initio* 3D reconstructions that were used for iterative rounds of heterogeneous refinements, where "junk" volumes generated from discarded particles served as decoy classes to capture and remove low-quality particles. For the scaffolded RNA-only datasets, 3D classification with a focus mask corresponding to the SLA from refined 3D reconstructions was used to resolve different SLA conformations, followed by signal subtraction of the scaffold and local refinement of the SLA density. For SLA-NS5 complexes, the final 3D reconstructions were refined using non-uniform and CTF refinement in cryoSPARC followed by 3D refinement using Blush regularization in RELION (*50*).

**Structural modeling**

*Modeling of DENV2, ZIKV, WNV, and YFV SLAs in scaffold constructs.* The structural modeling of the scaffold RNA constructs was performed using auto-DRRAFTER (Rosetta v. 3.14) (*41*). Composite maps of the locally refined SLA states and non-uniform refined map were used. Auto-DRRAFTER runs were setup manually by docking the TET and cpTET scaffold structures (PDB: 7EZ0) (*39*) into the map. A-form helices of SLA stems were generated using Rosetta's rna_helix.py script. One auto-DRRAFTER round with 5,000 decoys was run using the flags:

```
-secstruct_file $SECSTRUCT.TXT

-s $SCAFFOLD.PDB $H1.PDB $H2.PDB $H3.PDB $H4.PDB $H5.PDB

-edensity:mapfile $MAP
```

-edensity:mapreso $RESOLUTION

-out:file:silent $FILE

-nstruct 50

-cycles 1000

-dock_into_density false

-new_fold_tree_initializer true

-ft_close_chains false

-bps_moves false

-minimize_rna true

-minimize_protein_sc true

-rna_protein_docking true

-rnp_min_first true

-rnp_pack_first true

-rnp_high_res_cycles 2

-minimize_rounds 1

-ignore_zero_occupancy false

-convert_protein_CEN false

-FA_low_res_rnp_scoring true

-ramp_rnp_vdw true

-docking_move_size 0.5

-dock_each_chunk_per_chain false

-mute protocols.moves.RigidBodyMover

-mute protocols.rna.denovo.movers.RNA_HelixMover

-use_legacy_job_distributor true

-no_filters

-set_weights linear_chainbreak 20.0

-jump_library_file RNA18_HUB_2.154_2.5.jump

-vall_torsions RNA18_HUB_2.154_2.5.torsions

-score:weights stepwise/rna/rna_res_level_energy4.wts

-restore_talaris_behavior

-edensity:cryoem_scatterers

*Modeling of DENV2 and ZIKV SLA-NS5 complexes.* Starting models of DENV2 and ZIKV NS5 were generated by AlphaFold 3 (*75*) and rigid body fitting of the MTase and RdRp domains was performed in UCSF ChimeraX-1.9 (*63*). The median models of the S-T states of DENV2 and ZIKV generated from auto-DRRAFTER were docked into their respective SLA-NS5 map. Next, we performed molecular dynamics flexible fitting using the web server Namdinator (*76*) using the following parameters: Start & final temperature: 298 K, G-force scaling factor: 0.1, Minimization steps: 5,000, Simulation steps: 50,000, Phenix RSR cycles:1, Implicit solvent: exclude. Further refinement was performed using real-space refinement (RSR) in Phenix (v. 2.0). 100 max iterations with 5 macro cycles were run with restrained secondary structure and "minimization_global", "occupancy", "nqh_flips", "local_grid", and "adp". We then further refined the SLA using ERRASER2 (https://docs.rosettacommons.org/docs/latest/ERRASER2) using the following flags:

-s $PDB

-edensity:mapfile $MAP

-fasta $FASTA

-edensity:mapreso $RESOLUTION

-score:weights stepwise/rna/rna_res_level_energy7beta.

-set_weights elec_dens_fast 10.0 cart_bonded 5.0 linear_chainbreak 10.0 chainbreak 10.0 fa_rep 1.5 fa_intra_rep 0.5 rna_torsion 10 suiteness_bonus 5 rna_sugar_close 10

-mute core.scoring.CartesianBondedEnergy core.scoring.electron_density.xray_scattering

-rounds 3

-cryoem_scatterers

-rmsd_screen 3.0

-ignore_zero_occupancy false

-missing_density_to_jump

-allow_virtual_side_chains false

-pack_protein_side_chains false

Several iterative rounds of real space fitting with restraints in COOT (v. 0.9.8.96), RSR in Phenix (v. 2.0) and ERRASER2 were performed to reduce steric clashes and improve geometry of the model. Finally, the 5' cap was added to our models by grafting the cap from the capped DENV4 SLA bound to DENV3 NS5 model (PDB: 8GZP). A final round of Phenix RSR was performed with 50 max iterations and 1 macro cycle using the same parameters as described.

**smFRET sample preparation, data acquisition, and analysis**

Custom RNA oligos containing internal 5-aminohexylacrylamino-uridine modifications (Dharmacon Horizon; sequences and labeling position are shown in Table S2) were conjugated to mono reactive NHS ester Cy3 (PA13101, Cytiva) and Cy5 (PA15101, Cytiva). Following ethanol precipitation and 2' ACE deprotection, the Cy3/Cy5 labelled fragments were ligated using a 30-nucleotide DNA splint (15-nt overlap with each fragment; Table S2) and T4 RNA ligase 2 (M0239L, New England Biolabs). The ligated sample was treated with DNase I (M0303S, New England Biolabs), ethanol precipitated and purified by denaturing gel electrophoresis.

Folding of Cy3/Cy5-labeled SLA was evaluated by native polyacrylamide gel electrophoresis and EMSA. 20 nM RNA was refolded in 50 mM Tris-HCl pH 7.5 by heating to 80°C for 1 min and subsequently placing on ice. For EMSA, complex assembly was done as described above (see "Electromobility shift assays (EMSAs)" section). 2 µL of 50% (v/v) glycerol was added to the 10µL sample and the mixture was resolved on a 5% (w/v) polyacrylamide gel with 5% (v/v) glycerol.

Alternating laser excitation (ALEX)-based smFRET data was acquired on an EI-FLEX bench-top microscope (Exciting Instruments Ltd, Sheffield, UK). The Cy3/Cy5-labeled SLA was refolded in 50 mM Tris-HCl pH 7.5 by heating to 80°C for 1 min and subsequently placing on ice. 50 µL of refolded RNA sample equilibrated in buffer (50 mM Tris-HCl pH 7.5, 0-20 mM $MgCl_2$, 50 µg/mL BSA) was placed onto a no. 1 thickness coverslip and excited with alternating 520 nm (0.08 mW) laser and 638 nm (0.05 mW) lasers, respectively. Data were collected for 60 min at room temperature using 100 µs ALEX periods, with each laser alternating between 45 µs on and 5 µs off cycles.

Data analysis was performed with Jupyter Notebooks using the FRETBursts python module (*77*) adapted from smfBox pipeline (https://craggslab.github.io/smfBox/). As done previously (*78*), corrected FRET efficiencies of all doubly-labeled bursts were obtained after background subtraction from each channel and determination of correction factors. Correction factors ($\alpha$; leakage factor, $\delta$; direct acceptor factor, $\gamma$; detection factor, $\beta$; excitation factor) were determined for each set of replicates separately from a single measurement at 5 mM $MgCl_2$. The same correction factors were used for G17A SLA as for WT, as the dye environment is not expected to change as the mutation is distal to the labeling site. Dual channel burst search was performed using the following parameters; $m$ = 10, $f$ = 15, and $l$ = 20, where $m$ is the photon window size, $f$ represents the fold increase above background required to define a burst, and $l$ specifies the minimum photon count required for an event to be considered a burst. For WT SLA smFRET experiments with varying $MgCl_2$ concentration, FRET efficiency histograms were normalized and fitted to a two-Gaussian model in which sigma values were shared across replicates within each $MgCl_2$ concentration, while means and amplitudes were fitted independently for each replicate. Relative populations of low and high FRET states were calculated as the fractional area under each Gaussian component of the two-Gaussian fit.

**Bioinformatic analysis of flavivirus SLAs**

A covariance model-based bioinformatic analysis of SLAs from diverse flaviviruses was performed using Infernal v. 1.1.5 (*43*) and R-scape v. 2.6.4 (*79*).

*Generation of covariance model*. A sequence and secondary structure consensus model were built from a modified Rfam alignment (RF02340) that contained 17 sequences from DENV, WNV, Japanese encephalitis virus, and ZIKV, to which we added a YFV and tick-borne encephalitis virus sequence. The seed alignment and consensus secondary structure was adjusted using MAFFT (*80*) and RNAalifold (*81*), as well as making minor manual adjustments. The seed alignment was used to build and calibrate the covariance model.

*Search database curation, structural alignment and conservation analysis*. All complete flavivirus genome sequences were retrieved from National Center for Biotechnology Information (NCBI) virus database (last retrieved on 13 March 2026). To capture the 5' UTR comprising SLA, the sequences were trimmed to the first 200 nucleotides, duplicates were removed and filtered to retain those that contain the conserved 5' AG dinucleotide. The resulting search database containing 1157 sequences was queried using Infernal's cmsearch command using default parameters, resulting in 955 hits with E-values <0.01 from 35 flavivirus species. To reduce bias from over-represented flavivirus variants, a new sequence alignment using one representative SLA sequence per species was generated using Infernal. The consensus nucleotide sequence and secondary structure, as well as significant covariation of base pairing elements, were evaluated with R-scape with default parameters. SLA 3WJ logo plot was generated using WebLogo 3 webserver (https://weblogo.threeplusone.com/create.cgi).

**Native mass spectrometry**

2.5 μM DENV2 NS5 was mixed with SLA RNA at 1:1 molar ratio. The samples (NS5 only and NS5 with SLA RNA) were buffer-exchanged twice into 150 mM ammonium acetate pH 7.5, 0.01% (v/v) Tween-20 using Zeba microspin desalting columns with a 7-kDa molecular weight cutoff

(Thermo Scientific). For MS characterization, an aliquot of the buffer-exchanged sample was loaded into a gold-coated quartz capillary tip that was prepared in-house and then electrosprayed into an Exactive Plus with extended mass range (EMR) instrument (Thermo Fisher Scientific) with a static direct infusion nanospray source (*82*). The native MS parameters included spray voltage, 1.22 kV; capillary temperature, 150°C; in-source dissociation, 10 V; S-lens RF level, 200; resolving power, 8,750 at m/z of 200; AGC target, 1 x $10^6$; maximum injection time, 200 ms; number of microscans, 5; total number of scans, at least 100. Additional MS parameters were injection flatapole, 8 V; interflatapole, 7 V; bent flatapole, 4 – 6 V; high energy collision dissociation (HCD), 200 V; ultrahigh vacuum pressure, 4.2 × $10^{-10}$ mbar. Mass calibration in positive extended mass range mode was performed using cesium iodide. For data processing, the collected MS spectra were visualized using Thermo Xcalibur Qual Browser (v. 4.2.47). Spectral deconvolution was performed using UniDec v. 4.2.0 (*83, 84*) with the following general parameters: gaussian smoothing, 0 - 1; background subtract curved, 10; smooth charge state distribution, enabled; peak shape function, Gaussian; Degree of Softmax distribution (beta parameter): 10. The observed mass deviation from the MS measurements (calculated as the % difference between the measured and expected masses relative to the expected mass) ranged from 0.0003 – 0.057%.

**RNA motif search**

All PDB entries in v. 4.35 (9021 structures) of the FR3D RNA database were searched with WebFR3D (https://rna.bgsu.edu/fr3d/) using the constraints matrix in Table S3. These constraints enforce positions 1-3 of the motif to be consecutive, a chain break between positions 3 and 4, a sugar-Hoogsteen base-pair between positions 1 and 4 as well as stacking interactions between positions 2,3 and 4. This search returned 3227 hits. These results were filtered to remove duplicate entries referring to the same RNA but deposited multiple times in the PDB. This left 185 unique instances of the motif which are listed in Supplementary Data 1.

Sequence alignments from previously published work and Rfam were selected for the raiA element (*85*), the c-di-AMP riboswitch (Rfam 00379) and group II introns (Rfam 02001). To identify the columns in the alignment corresponding to the GRR/A TL-like motif, sequences of PDB structures of raiA (*86*), c-di-AMP riboswitch (*64*) and group II introns (*87*) were aligned to the sequence alignments using Infernal v. 1.1.5 (*88*). Sequence alignments were curated to contain only sequences where N1-N3 positions were contiguous (no insertions) and where all four residues (N1-N4) were present. This resulted in the inclusion of 1379/2377 sequences for raiA, 102/107 sequences for the c-di-amp riboswitch and 405/407 sequences for the group II introns. Logo plots of GRR/A TL-like motifs were generated using WebLogo 3 webserver (https://weblogo.threeplusone.com/create.cgi).

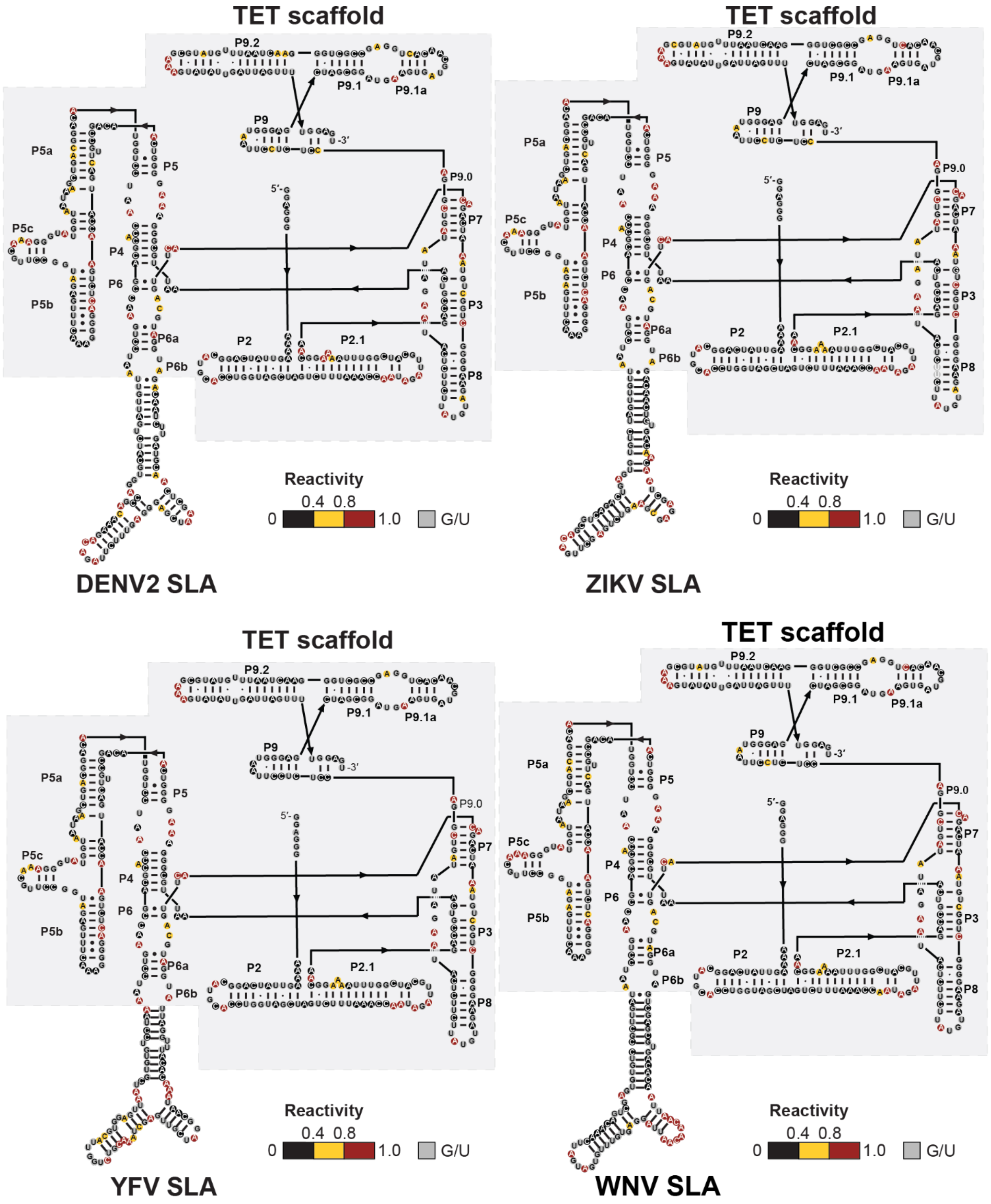

**fig. S1. DMS-MaPseq of SLA-TET constructs.** Normalized DMS reactivities are overlaid with consensus secondary structures for TET and SLAs from DENV2, ZIKV, YFV, and WNV. Low (0-0.4; black), medium (0.4-0.8; yellow) and high (0.8-1.0; red) reactivities are shown. G and U cannot be probed by DMS and are shown in grey.

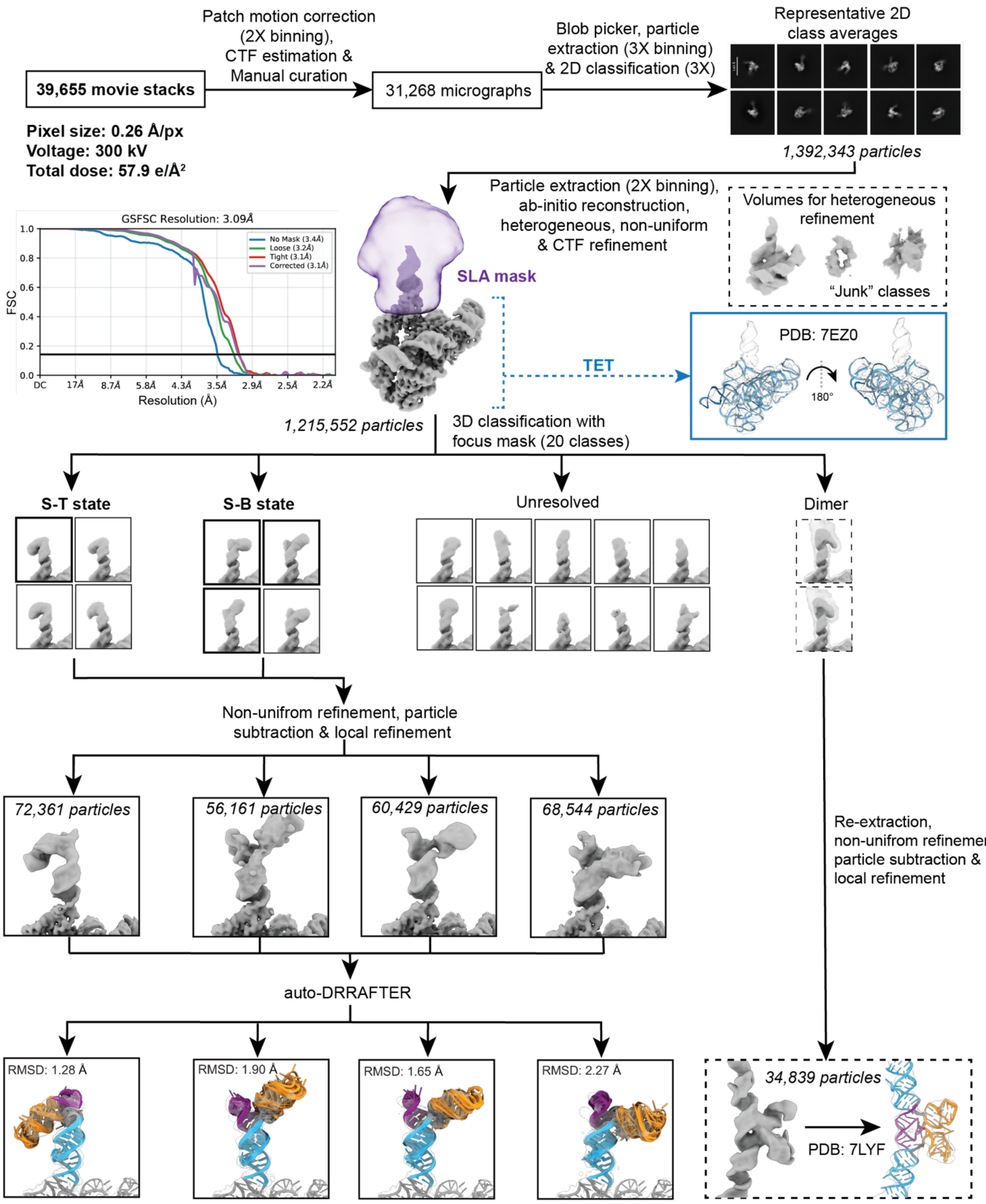

**fig. S2. Cryo-EM analysis of DENV2 SLA-TET.** Cryo-EM data analysis was performed with cryoSPARC (*72*). Three rounds of 2D classification were performed to remove "junk" particles. After 2D classification, particles were used for *ab initio* 3D reconstruction, which produced a volume consistent with the known 3D structure of TET (*39*). In addition to this volume, a set of junk particles was used to generate two 3D reconstructions that were used as "sinks" for a series of heterogeneous refinements. The final set of particles (n = 1,215,552) was used for non-uniform and CTF refinements, yielding a 3.1 Å resolution map. In this map, the density corresponding to the SLA was not well-defined. A mask at the SLA region was generated (shown in purple) and used for focused 3D classification into 20 classes. Classes that had densities consistent with a complete SLA—i.e., densities where the three stems of SLA were visible—were further refined after subtraction of the density corresponding to the TET scaffold. The composite maps, containing the refined SLA and TET densities were used for structural modeling using auto-DRRAFTER (*40*). The ten top-scoring auto-DRRAFTER models generated for each state are shown in cartoon representation using the color scheme from Fig. 1A, along with their mean pairwise RMSD. Densities consistent with dimers were also observed during 3D classification. These particles were re-extracted using a larger box and refined using non-uniform and focused refinement. The density matches the previously determined crystal structure of a DENV2 SLA dimer (*35*).

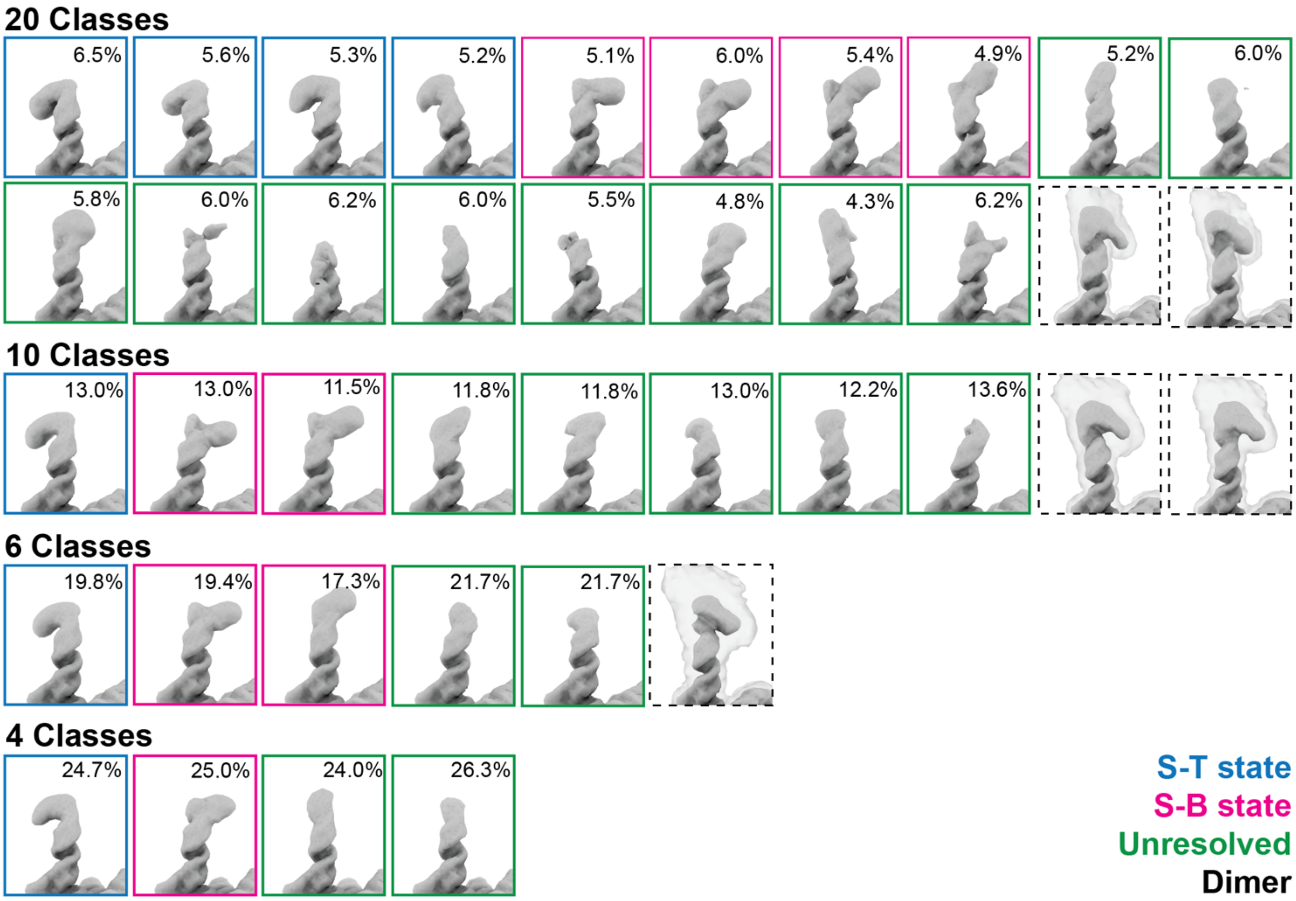

**fig. S3. Analysis of DENV2 SLA-TET 3D classes.** Particles that yielded the 3D reconstruction solved to 3.1 Å in fig. S2 were used for focused 3D classification of the SLA conformations, using a variable number of 3D classes. Classes that displayed density consistent with a complete SLA—i.e., where the three stems were visible—were classified as S-T state (blue) or S-B state (pink), dependent on their apparent coaxial stacking configuration and the relative orientations of their helices. Classes with densities that were not consistent with a complete SLA were classified as unresolved (green box). Classes with densities that extended beyond the size of an SLA were classified as dimers (black dashed box). Percentages were calculated for particles in different monomeric states.

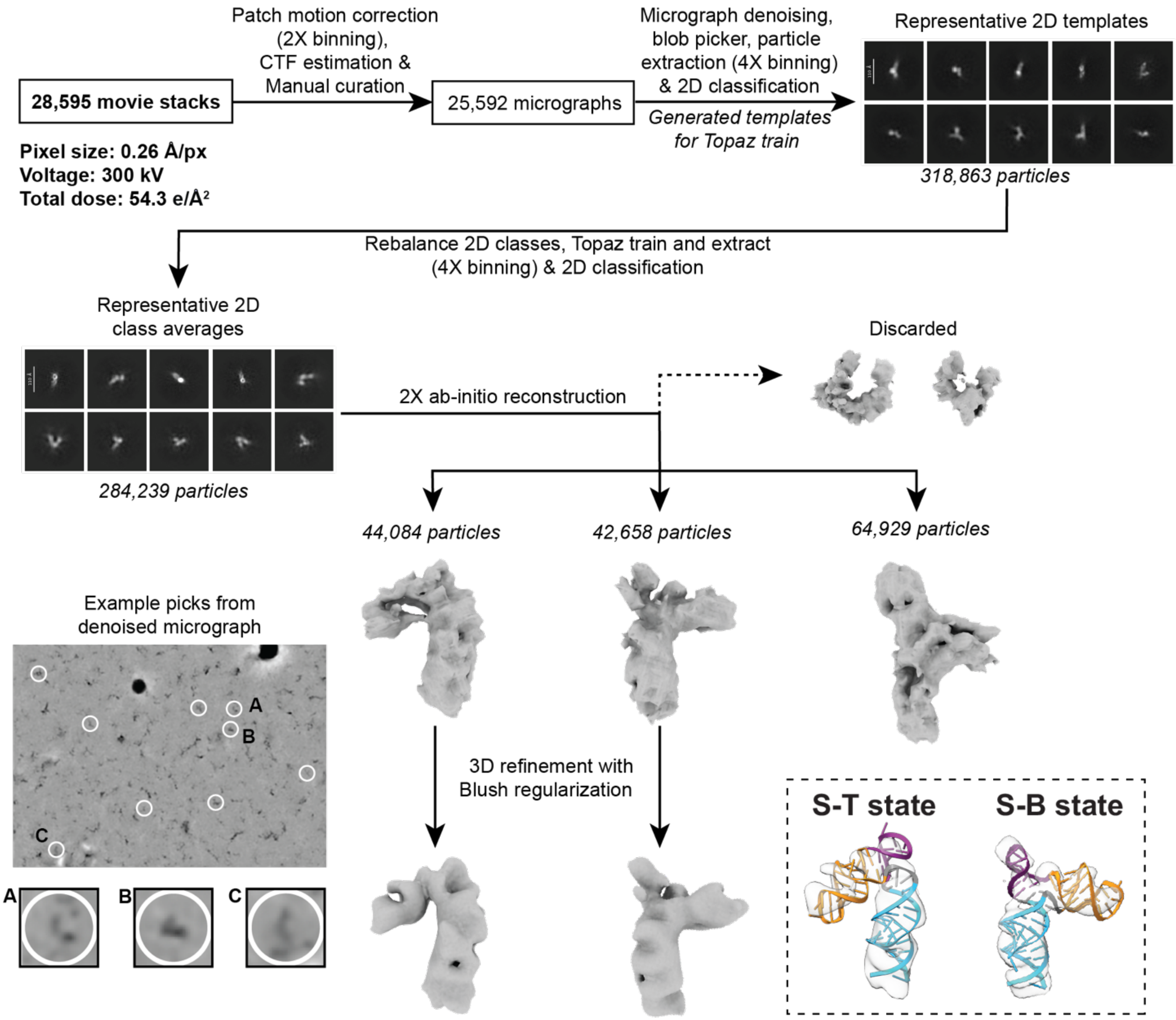

**fig. S4. Cryo-EM analysis of 70-nt DENV2 SLA construct.** Initial data analysis was done in cryoSPARC (*72*). Initial particle picking was done using blob picker. Due to the small size of the particles, multiple rounds of 2D classification were performed to generate templates for Topaz picking, a method that uses convolutional neural networks to help retrieve challenging particles with high accuracy (*74*). Prior to Topaz training, 2D classes were rebalanced to remove over-represented views. Several rounds of 2D classification were performed after Topaz picking, followed by two rounds of *ab initio* 3D reconstruction. Particles that generated *ab initio* reconstructions inconsistent with the expected size of the SLA were removed. A final *ab initio* 3D reconstruction with three classes generated two volumes were consistent with the expected size and secondary structure of the SLA. These classes were refined with Blush regularization in RELION (*50, 73*). Examples of picked particles from a denoised micrograph are shown on the bottom left. In the bottom right box, the 3D structures of the S-T and S-B states generated from the scaffold-based cryo-EM structures (fig. S2) were docked into the two final maps generated from the 70-nt DENV2 SLA construct using the color scheme from Fig. 1A.

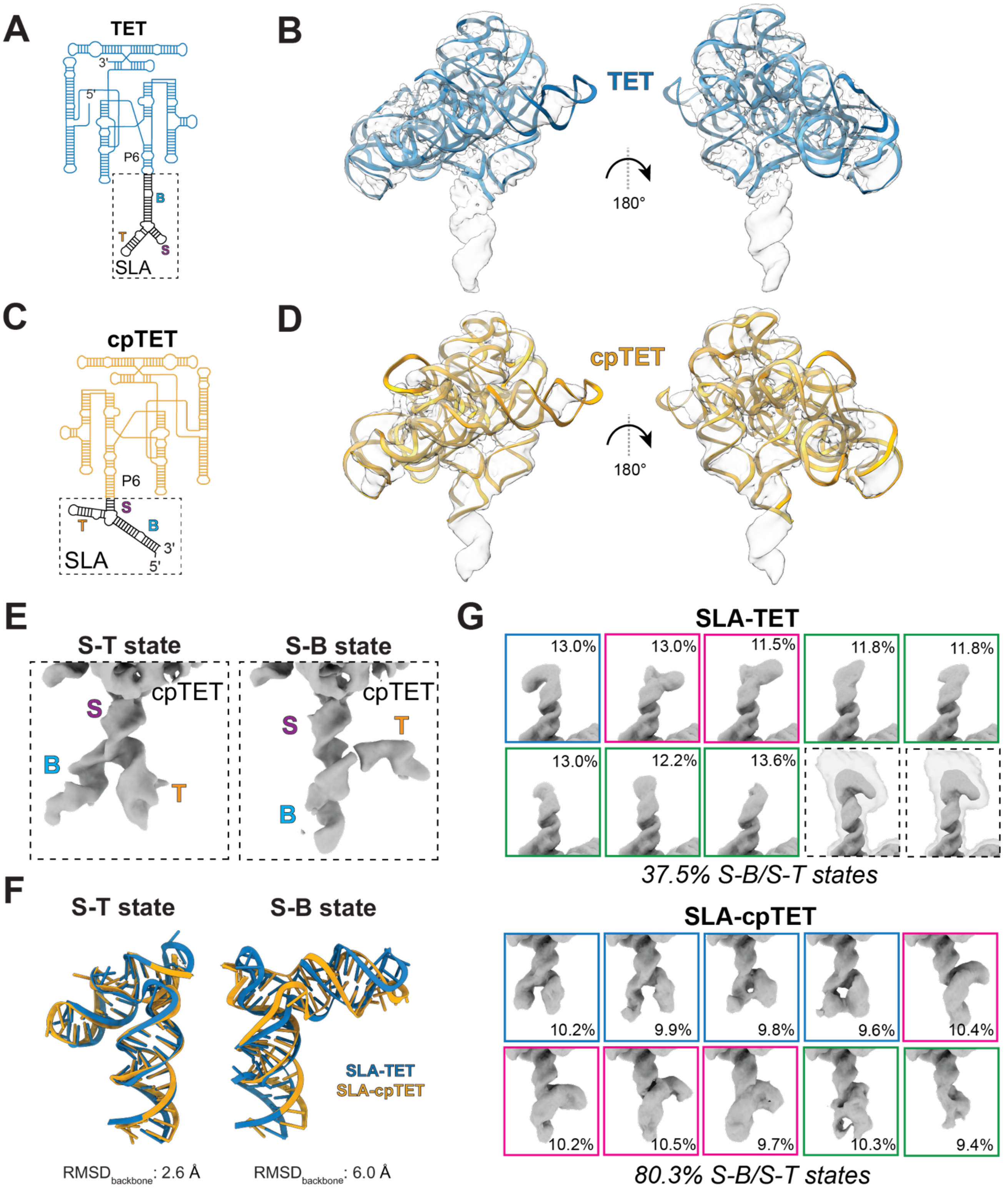

**fig. S5. DENV2 SLA appended to two different TET-based scaffolds.** (**A**) Secondary structure diagram of SLA appended to TET. The B stem of SLA was appended to the P6 helix of TET. (**B**) The 3D structure of TET (blue cartoon; PDB: 7EZ0) (*39*) is docked into the refined 3D density map of the TET-SLA construct. The 3D density map shown is that prior to 3D classification of the SLA. (**C**) Secondary structure diagram of SLA appended to a circularly permuted version of TET (cpTET). Note that the positions of the 5′ and 3′ ends differ from those of the original construct. The S stem of SLA was appended to the P6b helix of cpTET. (**D**) The 3D structure of TET (yellow cartoon; PDB: 7EZ0) (*39*) is docked into the refined 3D density map of the cpTET-SLA construct. The 3D density map shown is that prior to 3D classification of the SLA. (**E**) Focused 3D classification and refinement of DENV2 SLA in the cpTET scaffold construct produced two major conformational states. B, T and

S stems are labeled using the color scheme in Fig. 1A. Full cryo-EM data analysis workflow is shown below in fig. S6. (**F**) Superposition of 3D models obtained for the S-T and S-B states using the two different scaffolds. The structures shown are the median-scored models from auto-DRRAFTER modeling. The relatively larger $RMSD_{backbone}$ for the S-B state likely reflects the flexibility of this state, as analysis reveals multiple substates that are consistent with this coaxial stacking configuration (fig. S2). (**G**) Focused 3D classification of SLA in TET (top) and cpTET (bottom). Classes that displayed density consistent with a complete SLA—i.e., where the three stems were visible—were classified as S-T (blue box) or S-B (pink box) states, dependent on their apparent coaxial stacking configuration and the relative orientations of their helices. Classes with densities that were not consistent with a complete SLA were classified as unresolved (green box). Classes with densities that extended beyond the size of an SLA were classified as dimers (black dashed box). Percentages were calculated for particles in different monomeric states.

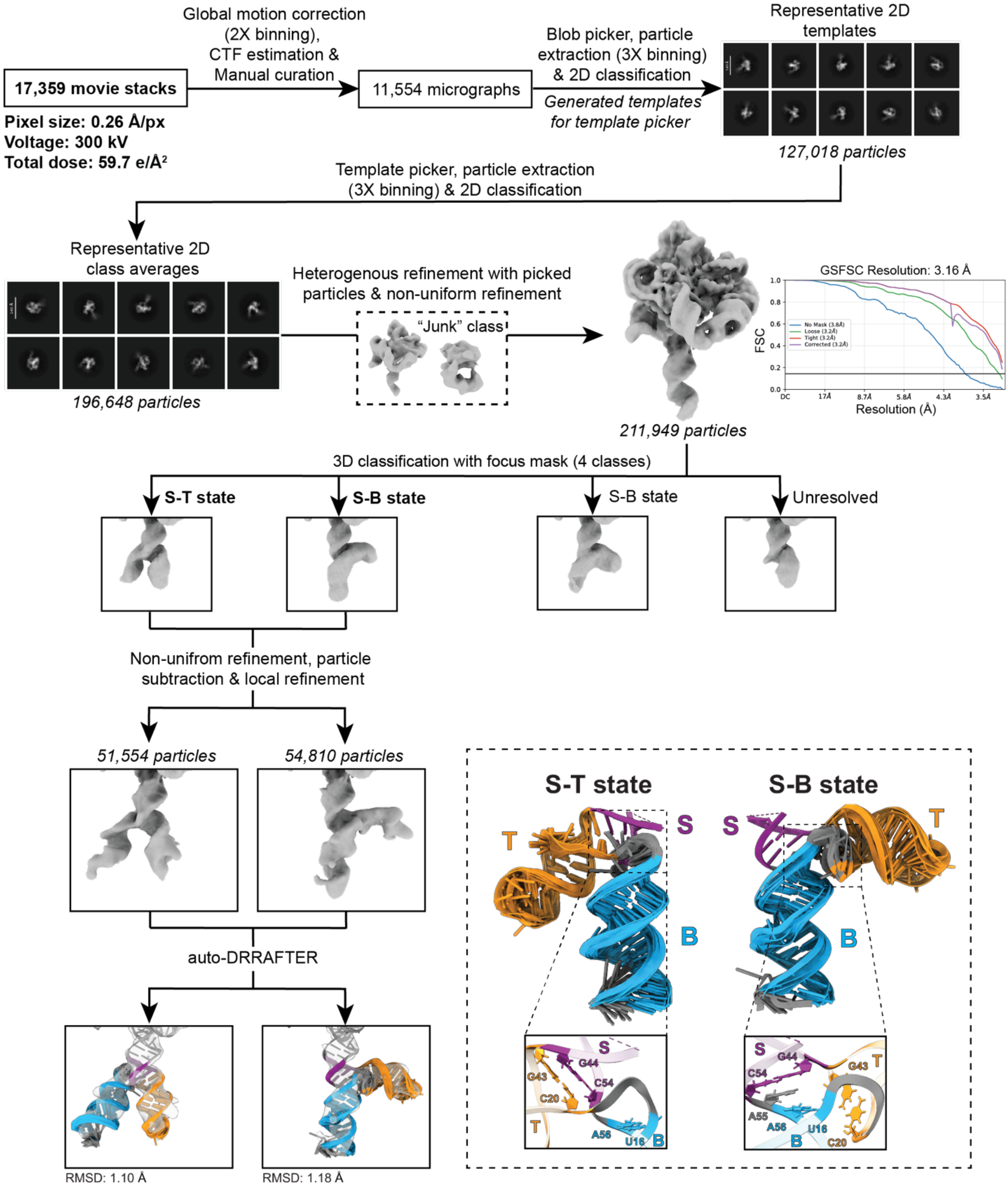

**fig. S6. Cryo-EM analysis of DENV2 SLA appended to cpTET scaffold.** Cryo-EM data analysis was performed with cryoSPARC (*72*). Particles were initially picked using blob picker and several rounds of 2D classification were performed to generate 2D templates for template picker. After several rounds of 2D classification to remove junk particles, particles were used for *ab initio* reconstruction, producing a volume that was consistent with the previously determined 3D structure of TET. As described above (fig. S2), heterogeneous refinement was used to classify particles and remove junk using a "sink" volume. Three rounds of heterogeneous refinement were performed. The final set of particles was used for non-uniform 3D refinements, producing a 3.2 Å resolution 3D reconstruction. The density corresponding to the SLA was not well-defined in this reconstruction. Focused 3D classification was performed using a mask around the SLA region. Two well-defined classes with densities that agreed with the size and secondary structure of SLA were refined using non-uniform and local 3D refinements. The composite maps were used for auto-DRRAFTER modeling (*40*); the ten top-scoring auto-DRRAFTER models are shown using the color scheme from Fig. 1A, along with their mean pairwise RMSD. Close-up view of the three-way junction (3WJ) of the two states show that they correspond to the same S-T and S-B states observed in the original scaffolded construct.

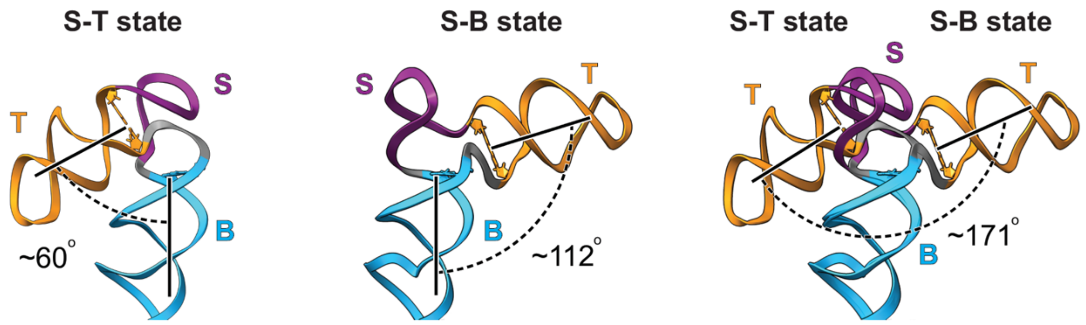

**fig. S7. Relative orientation of the B and T stem in the S-B and S-T state.** The angle between B and T stems for S-T (left) and S-B (middle) states from the median-scored SLA-TET auto-DRRAFTER models were calculated using vectors orthogonal to their closing base pairs. Comparison of the position of the T stem in S-T vs. S-B states revealed a rotation of ~171° (right). The two states were aligned by superimposing their B stem.

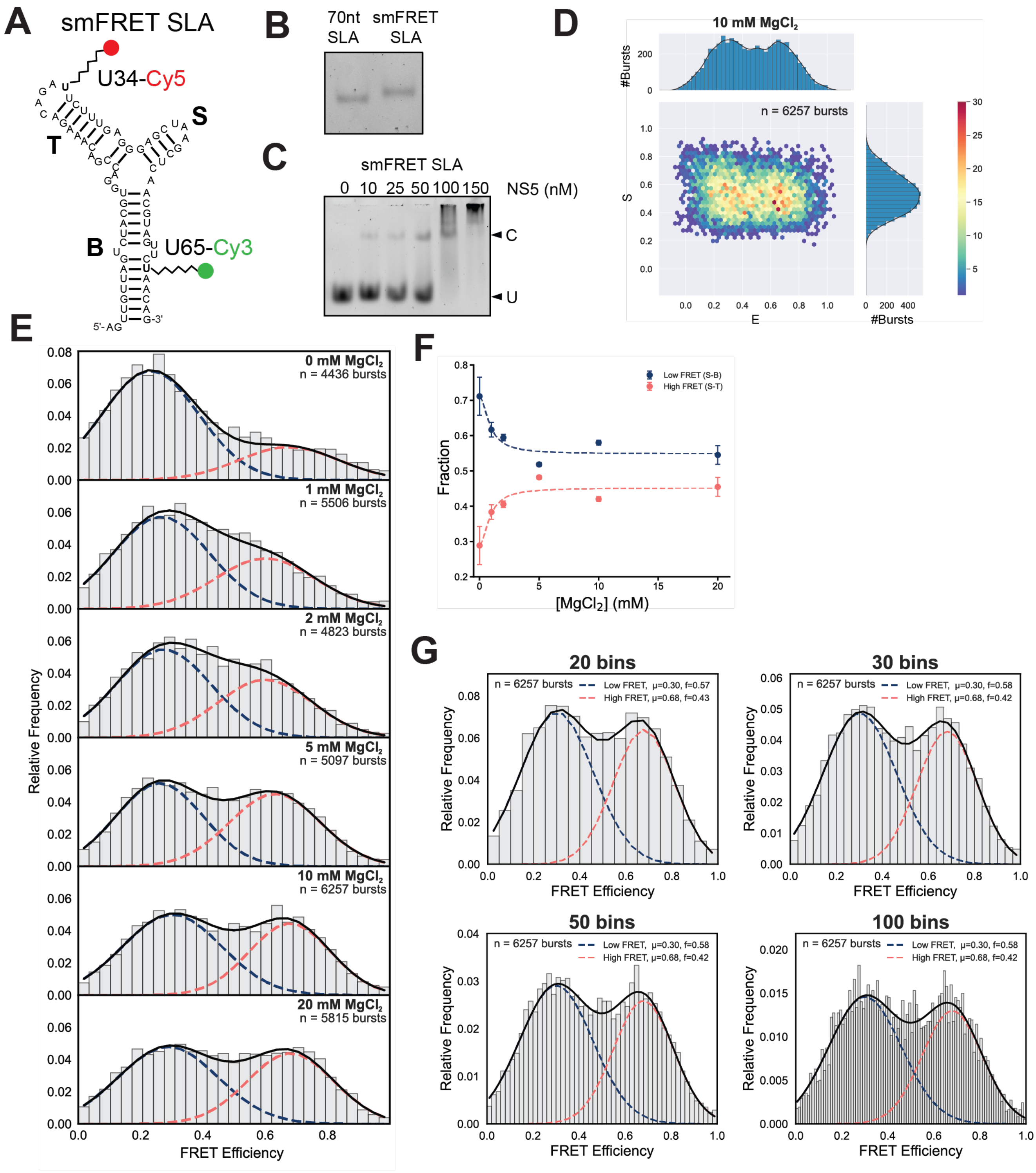

**fig. S8. smFRET studies of DENV2 SLA.** (**A**) DENV2 SLA labeling scheme for smFRET. Cy3 and Cy5 dyes were conjugated to positions U65 and U34, respectively, via a C6 linker. (**B**) Native 8% (w/v) polyacrylamide gel of refolded unlabeled and Cy3/Cy5-labeled SLA. (**C**) EMSA of Cy3/Cy5-labeled SLA with NS5 showing concentration-dependent binding. Bands corresponding to unbound SLA (U) and SLA-NS5 complex (C) are labeled. (**D**) Representative E vs S plot from dual-channel burst search showing selection of doubly-labeled bursts. (**E**) [$Mg^{2+}$]-dependent stabilization of high FRET state (S-T state). Normalized FRET efficiency histograms (30 bins) display bimodal distributions that were fit to a two-Gaussian model. Gaussian components corresponding to the low and high FRET states are shown in blue and pink, respectively. One representative replicate is shown. (**F**) Relative population of low (blue; S-B state) and high FRET (pink; S-T state) states as a function of [$Mg^{2+}$]. Points and error bars represent the mean and standard deviation of three replicates. Dashed lines show fits to the Hill equation. (**G**) Two-Gaussian model fitting with different bin sizes (20, 30, 50 and 100 bins). $\mu$ denotes the mean FRET efficiency and $f$ the relative population of each state, demonstrating bin-size independence of the fitted parameters.

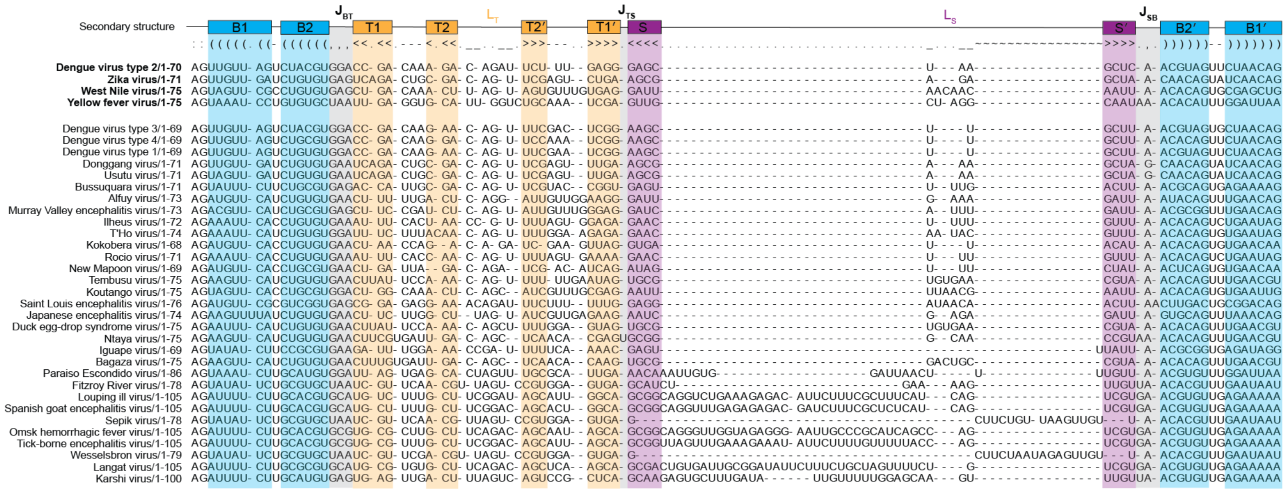

**fig. S9. Comparative sequence alignment of SLAs from diverse flaviviruses.** Infernal sequence alignment of SLAs from 35 diverse flaviviruses. The consensus secondary structure and sequences corresponding to SLAs from DENV2, ZIKV, WNV and YFV studied here are shown at the top. Regions corresponding to B (B1 and B2), T (T1 and T2), and S stems are labeled using same colors scheme as in Fig. 1A.

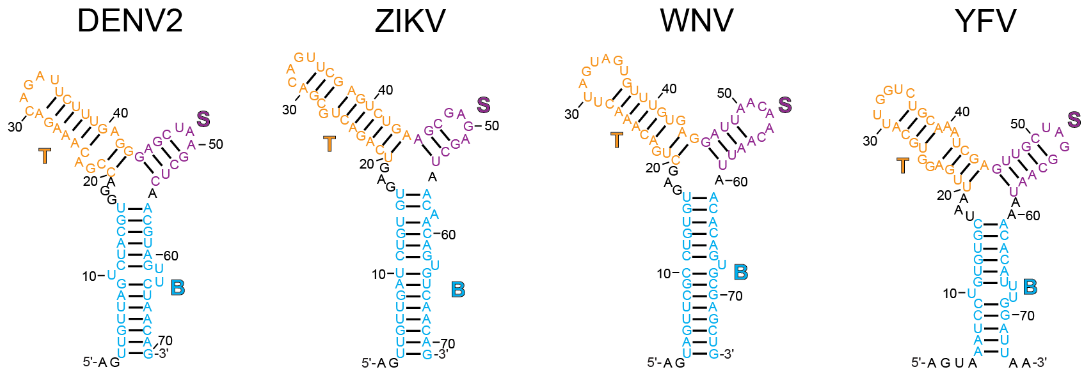

**fig. S10. Sequence and secondary structures of SLAs used.** Sequences and consensus secondary structures are shown, with B, T and S stems colored as in Fig. 1A.

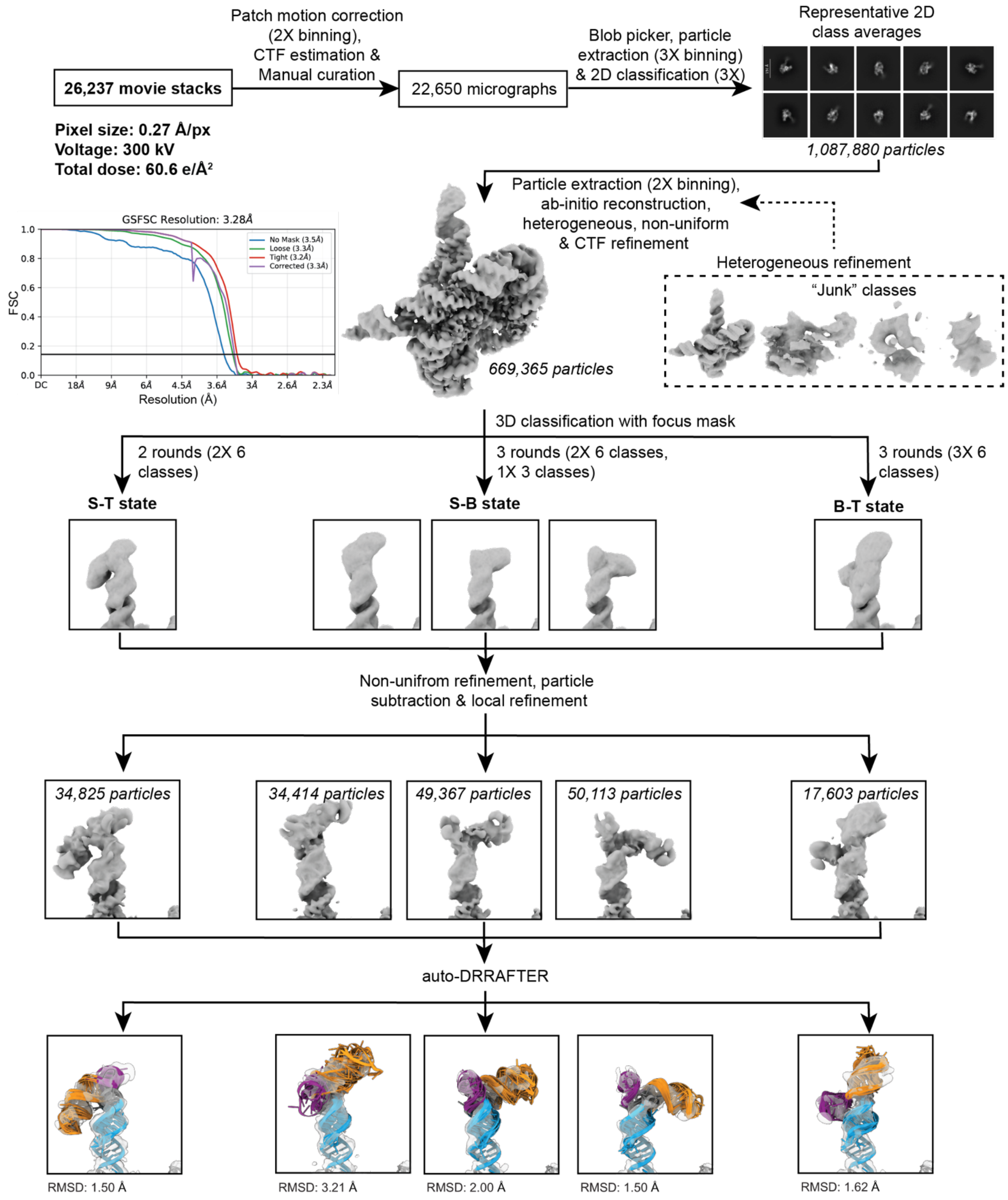

**fig. S11. Cryo-EM analysis of ZIKV SLA-TET.** Cryo-EM data analysis was performed with cryoSPARC (*72*). Particles were picked using blob picker and several rounds of 2D classification were used to remove junk particles and to generate *ab initio* reconstructions, producing a volume that was consistent with the previously determined 3D structure of TET. As described previously (fig. S2), heterogeneous refinement was used to classify particles and remove junk using a "sink" volume. Five rounds of heterogeneous refinements were performed. The final set of particles were used for non-uniform and CTF 3D refinements, producing a 3.3 Å resolution 3D reconstruction. Focused 3D classification was performed using a mask around the SLA region, using 6 classes. Classes that had densities consistent with a complete SLA were further 3D classified with a focus mask to resolve local structural fluctuations. The most complete and well-resolved 3D classes were further refined after subtraction of the density corresponding to the TET scaffold. One class with density consistent with the stacking of the B and T stem (B-T state) was observed. The composite maps, containing the refined SLA and TET densities were used for structural modeling using auto-DRRAFTER (*40*). The ten top-scoring auto-DRRAFTER models generated for each state are shown using the color scheme from fig. S10, along with their mean pairwise RMSD. The auto-DRRAFTER models confirmed the presence of the B-T state, indicating that all three possible stacking configurations of the 3WJ can be sampled.

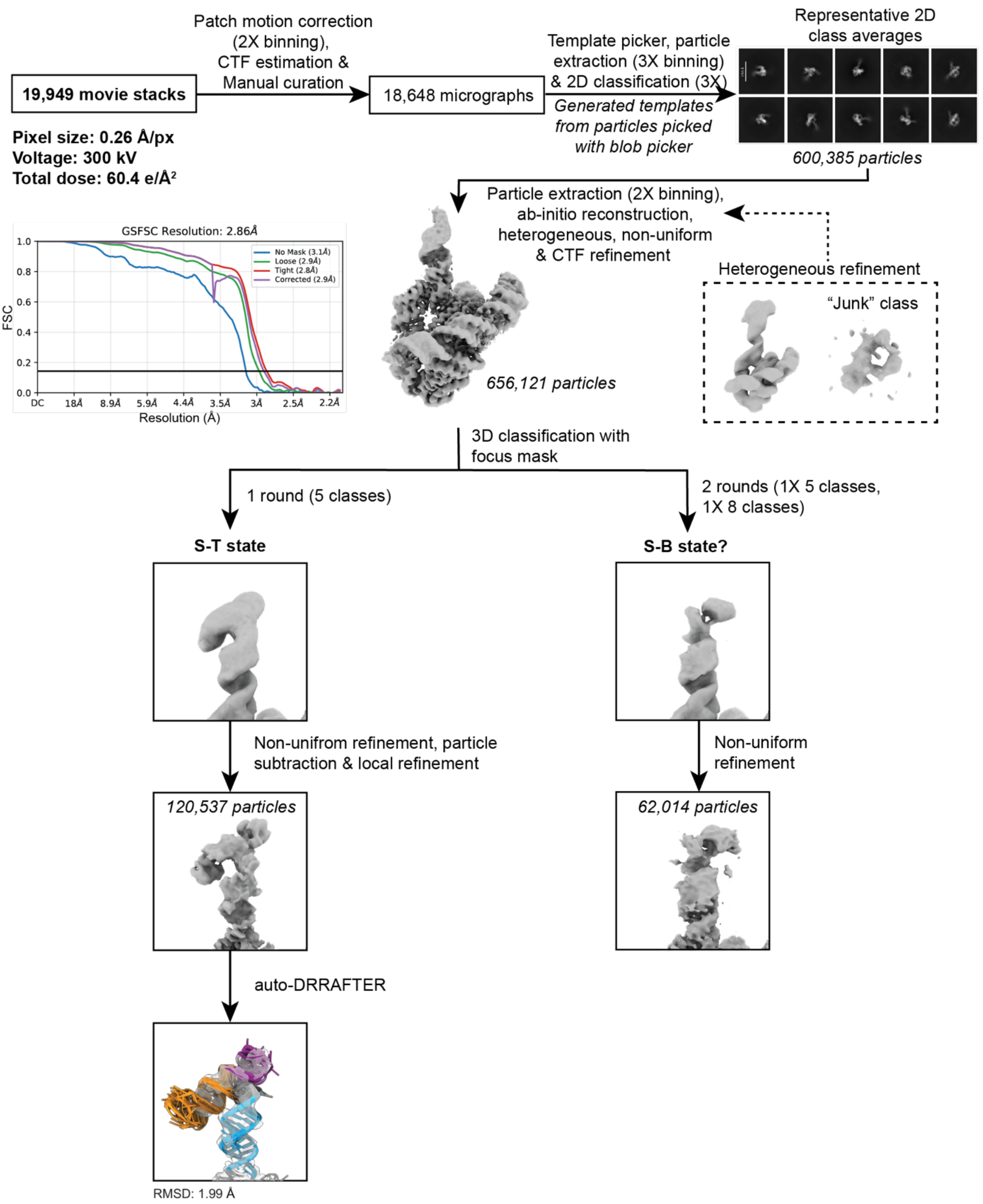

**fig. S12. Cryo-EM analysis of WNV SLA-TET.** Cryo-EM data analysis was performed with cryoSPARC (*72*). Multiple rounds of 2D classification were used to generate 2D templates for template picker. After several rounds of 2D classification to remove junk particles, particles were used for *ab initio* reconstruction and–as described above (fig. S2)–heterogeneous refinement was used to classify and remove junk particles using a "sink" volume. Five rounds of heterogeneous refinements were performed. The final set of particles was used for non-uniform and CTF refinements, producing a 3D reconstruction with an overall resolution of 2.9 Å. As done previously (fig. S2), focused 3D classification was performed using a mask around the SLA region, using 5 classes. One class resembled the S-T state, while the remaining particles were further classified in an additional round of masked 3D classification, using 8 classes. One of the densities resembled the S-B state but was insufficiently resolved to permit structural modeling. The composite map of the S-T state was used for structural modeling using auto-DRRAFTER (*40*). The ten top-scoring auto-DRRAFTER models generated for the S-T state are shown using color scheme from fig. S10, along with the mean pairwise RMSD.

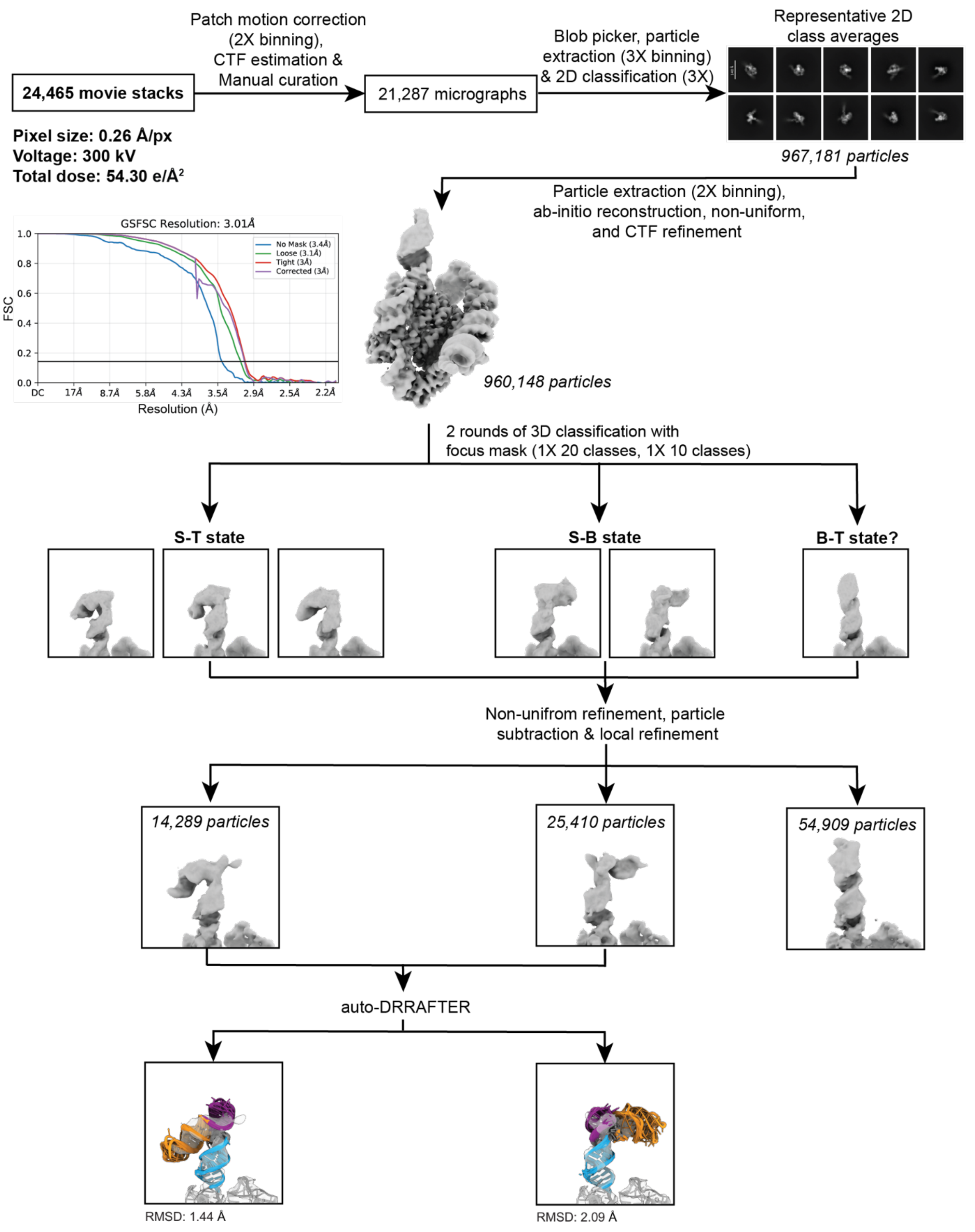

**fig. S13. Cryo-EM analysis of YFV SLA-TET.** Cryo-EM data analysis was performed with cryoSPARC (*72*). Particles were picked using blob picker and several rounds of 2D classification were used to remove junk particles and to generate *ab initio* reconstructions, producing a volume that was consistent with the previously determined 3D structure of TET. The 3D volume was refined using non-uniform and CTF 3D refinement, generating a 3.0 Å resolution 3D reconstruction. Masked 3D classification using 20 classes resulted in 3D classes resembling S-T, S-B and B-T states. As done previously (fig. S11), an additional round of masked 3D classification using 10 classes for each state was performed separately, further improving the resolution of each state. The particles from the best-defined classes in each state were combined, as they were very similar, and further refined after subtracting the density corresponding to the TET scaffold. The composite maps of the S-T and S-B states, containing the refined SLA and TET densities, were used for structural modeling using auto-DRRAFTER (*40*); the 3D class consistent with the B-T state was missing density to permit structural modeling of the S stem and was not modeled. The ten top-scoring auto-DRRAFTER models generated for each state are shown using the color scheme from fig. S10, along with their mean pairwise RMSD.

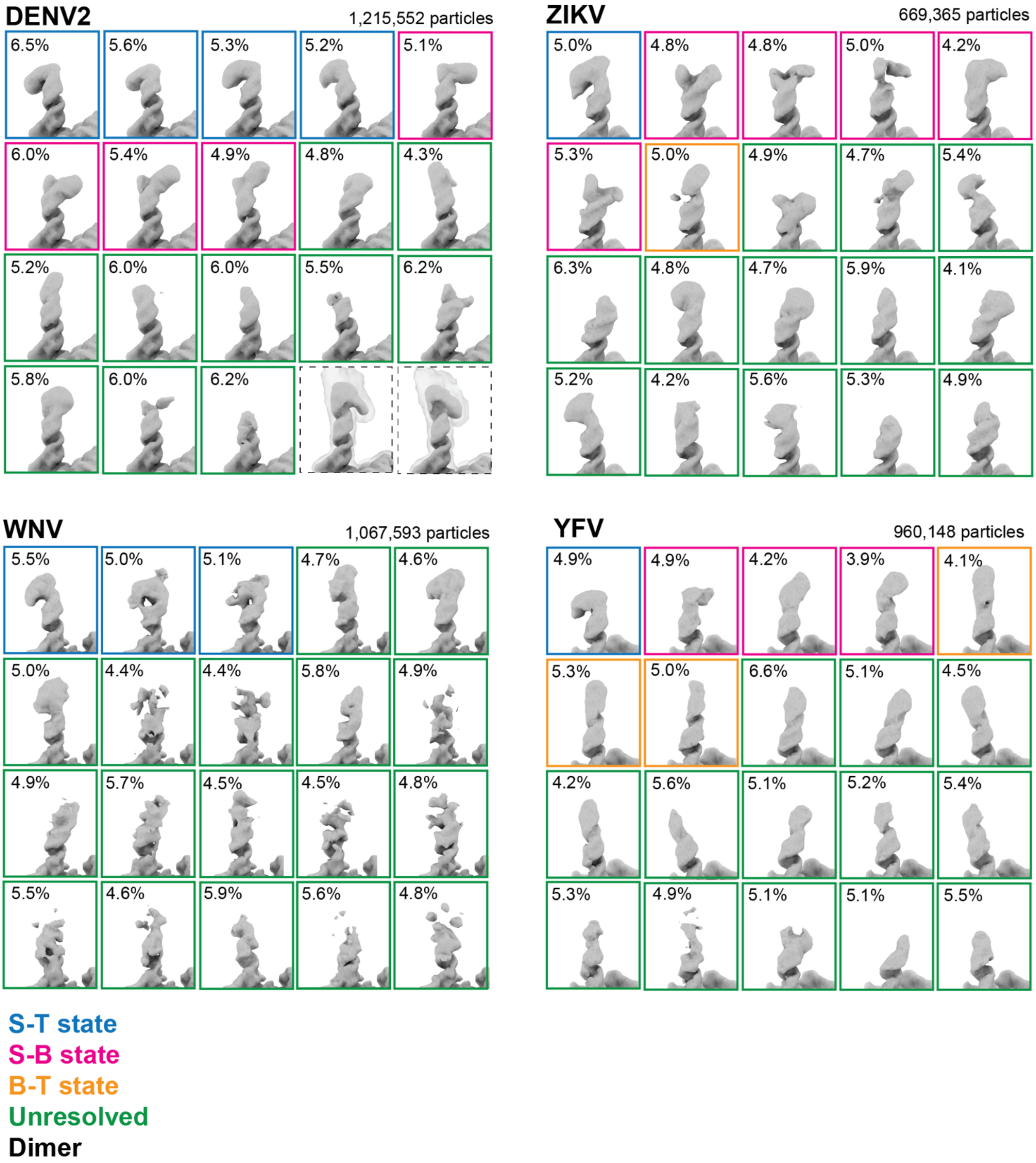

**fig. S14. Conformational heterogeneity across different flavivirus SLAs.** Focused 3D classification using a mask around the SLA region (as shown in fig. S2) with 20 classes was performed for DENV2, ZIKV, WNV and YFV SLA-TET datasets. Classes that displayed density with apparent coaxial stacking, based on the relative orientation of the helices, were classified as S-T state (blue), S-B state (pink), and B-T state (orange). Classes that that appeared incomplete were classified as unresolved (green). Classes with densities consistent with dimers are shown in black. The total number of particles used for each 3D classification is indicated for each dataset (top right) and particle percentages are shown for each class (calculated for particles in different monomeric states).

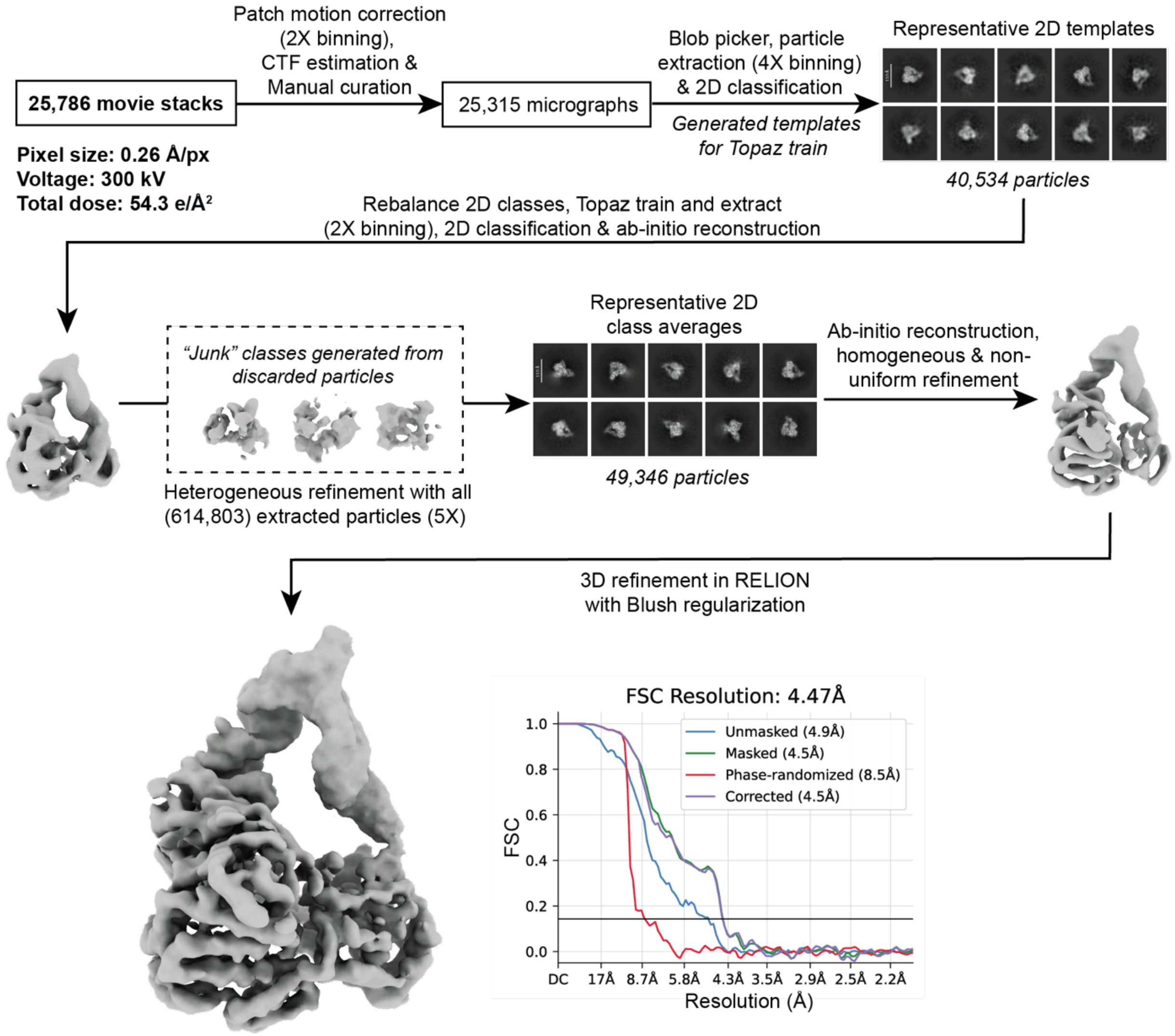

**fig. S15. Cryo-EM analysis of DENV2 SLA-NS5.** Initial data analysis was done in cryoSPARC (*72*). As described previously (fig. S4), multiple rounds of 2D classification and 2D rebalancing was performed to train Topaz picking model, using 50 epochs (*74*). Particles were picked with Topaz picker and multiple rounds of 2D classification to remove junk particles were used for *ab initio* reconstruction, producing a 3D volume consistent with the size of the SLA-NS5 complex. As described previously (fig. S2)–heterogeneous refinement was used to classify particles and remove junk using a "sink" volume. Five rounds of heterogeneous refinements were performed and followed by homogeneous and non-uniform refinements. A final round of 3D refinements using Blush regularization was performed in RELION (*50, 73*), resulting in a 4.5 Å resolution 3D reconstruction.

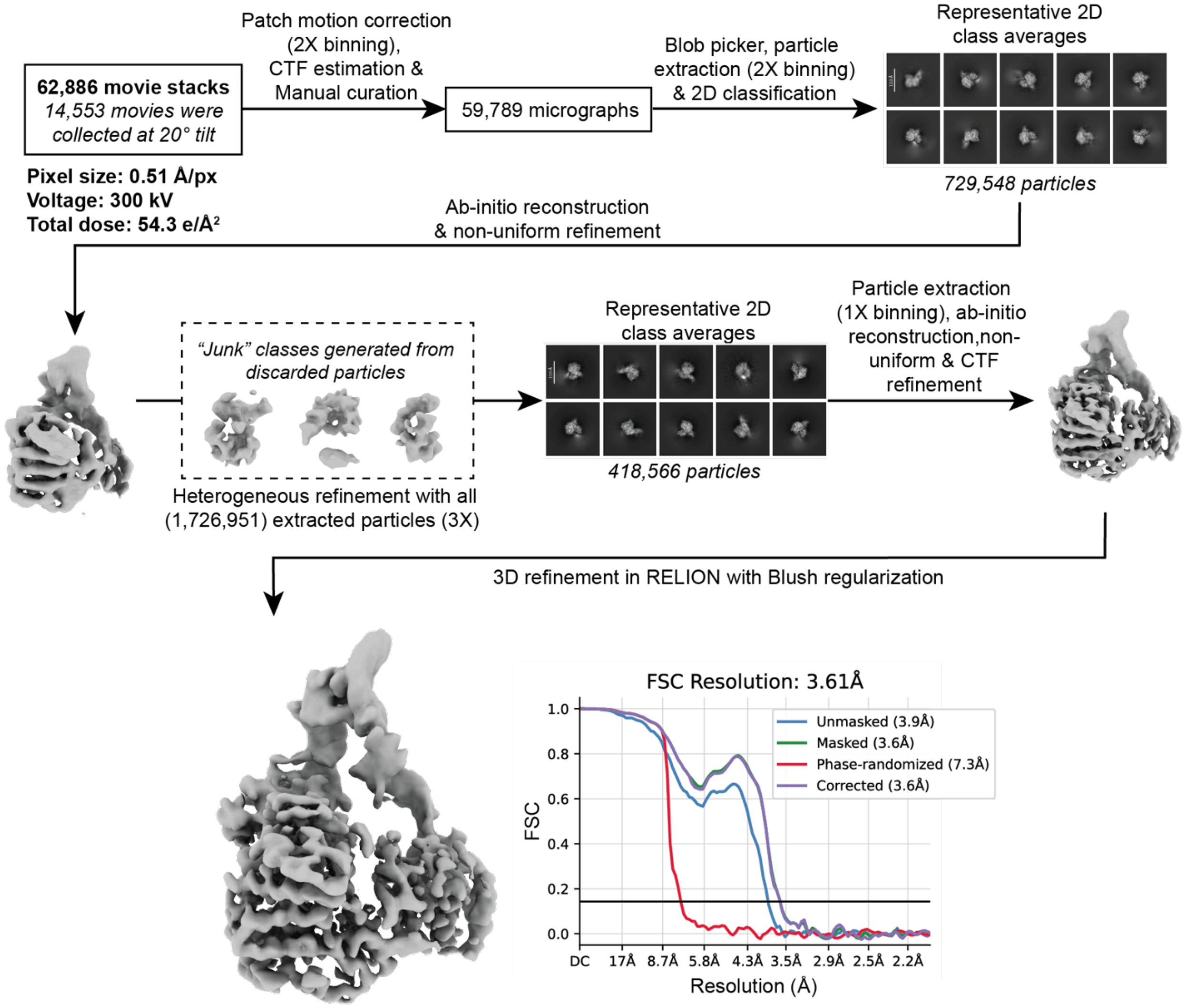

**fig. S16. Cryo-EM analysis of ZIKV SLA-NS5.** Initial data analysis was done in cryoSPARC (*72*). Particles were picked using blob picker and several rounds of 2D classification were used to remove junk particles and to generate *ab initio* reconstructions, producing a volume that was consistent with the expected size of the SLA-NS5 complex. As described previously (fig. S2), heterogeneous refinement was used to classify particles and remove junk using a "sink" volume. Three rounds of heterogeneous refinements were performed and followed by non-uniform and CTF refinements. A final round of 3D refinements using Blush regularization was performed in RELION (*50, 73*), resulting in a 3.6 Å resolution 3D reconstruction.

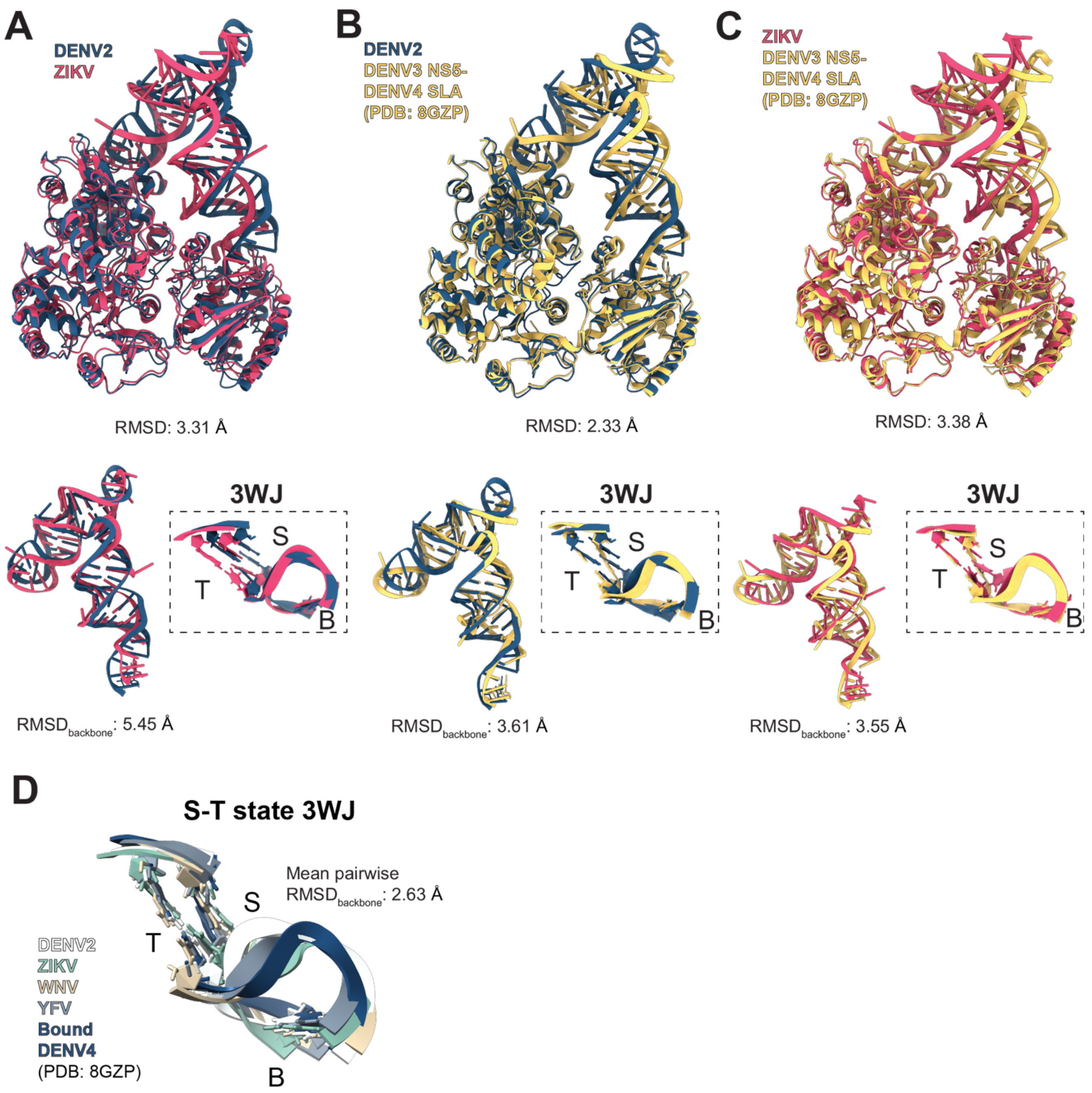

**fig. S17. Conserved NS5 recognition of SLA across flavivirus species and serotypes.** (**A-C**) Structural comparisons of DENV2 vs. ZIKV (**A**), DENV2 vs. DENV4 SLA-DENV3-NS5 (PDB: 8GZP) (*51*) (**B**) and ZIKV vs. DENV4 SLA-DENV3 NS5 (PDB: 8GZP) (*51*) (**C**) complexes. For each comparison, alignments of the full complex (top), SLA alone (bottom left) and 3WJ (bottom right) are shown. RMSD of the complex was calculated using the SLA phosphate backbone and NS5 $C_\alpha$ atoms in UCSF ChimeraX (*63*). SLA RMSD$_{backbone}$ was calculated over manually selected base-paired residues. (**D**) Median-scored auto-DRRAFTER models of the S-T states from unbound ensembles of DENV2, ZIKV, WNV and YFV superimposed onto NS5-bound DENV4 SLA (PDB: 8GZP) (*51*). RMSD$_{backbone}$ was calculated over residues of 3WJ including the closing base pairs.

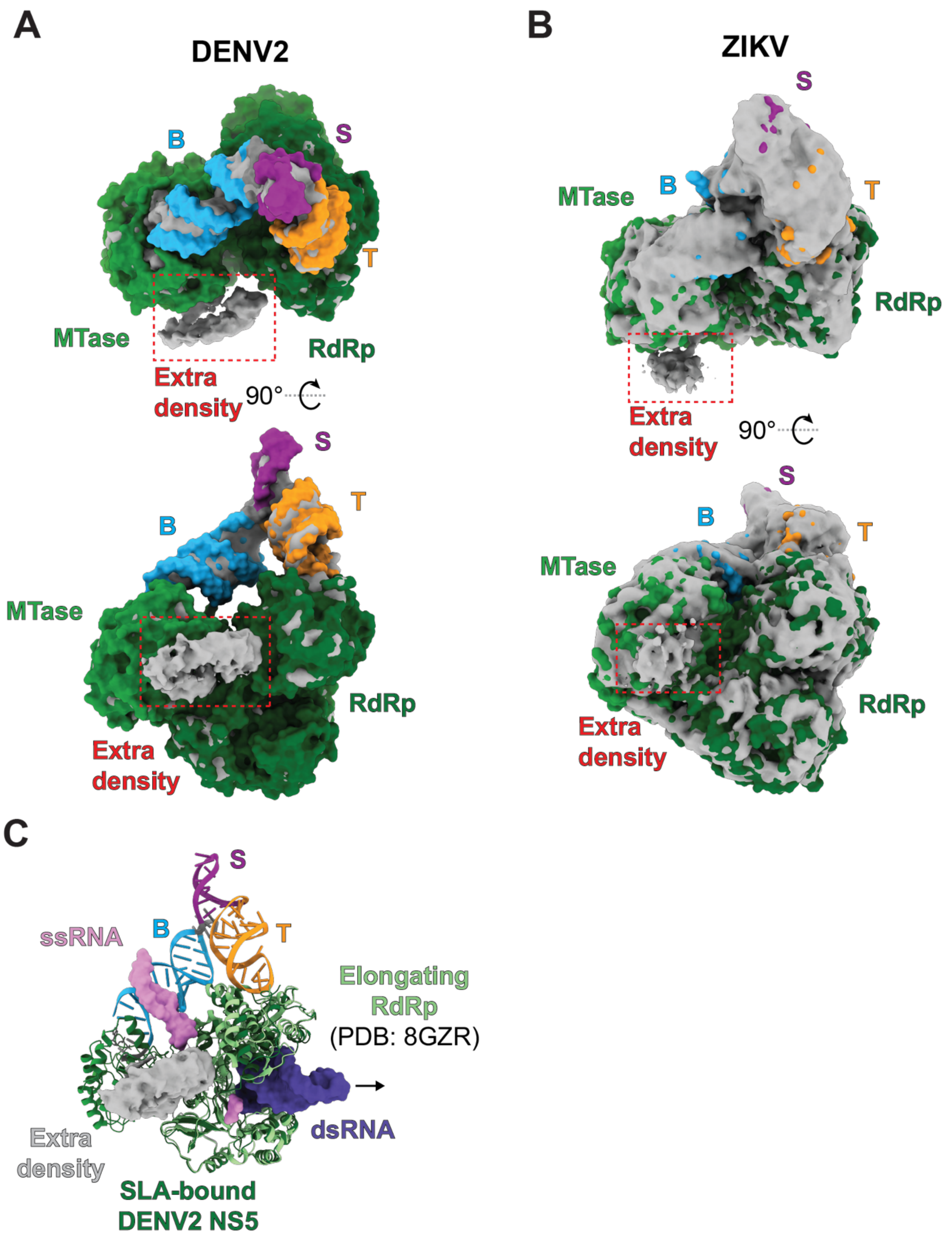

**fig. S18. Extra density observed in SLA-NS5 cryo-EM maps.** (**A**) Extra density near the RNA template entry site in the DENV2 SLA-NS5 cryo-EM map. The model, shown in surface representation, is docked into the cryo-EM density map at a low contour threshold, where extra density (outlined in red) is visible. (**B**) Extra density observed in the ZIKV SLA-NS5 cryo-EM map, shown using the same representation and coloring scheme as in panel A. (**C**) Superposition of the RdRp domain of the DENV2 SLA-NS5 complex onto the RdRp domain of an elongating DENV3 NS5-NS3 complex. The DENV2-SLA NS5 is shown in cartoon representation using same color scheme as in panel A. The elongating RdRp (lime green), incoming ssRNA template (pink) and dsRNA product (purple) are shown. The MTase domain of the elongating DENV3 NS5 and NS3 are not shown.

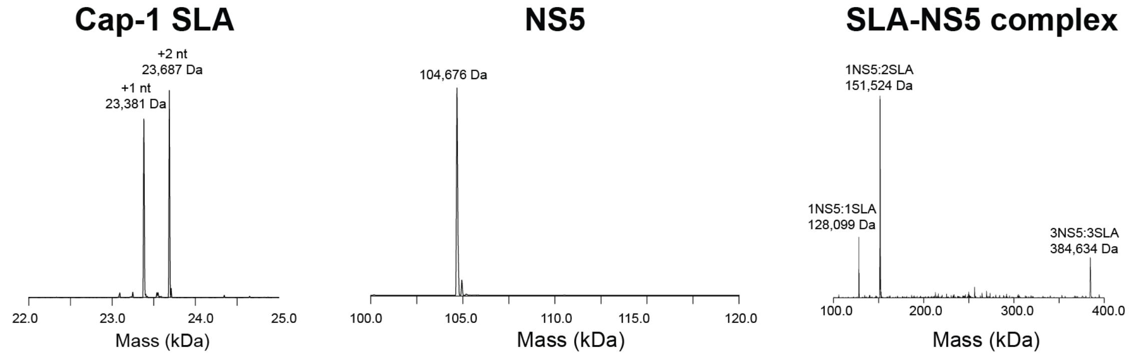

**fig. S19. Native mass spectra of SLA, NS5 and DENV2 SLA-NS5 complex.** Deconvolved MS spectra of 5′-capped SLA (left), NS5 (middle), and SLA-NS5 complex (right) are shown. Two major peaks are observed for cap-1 SLA, consistent with +1 and +2 nt non-templated nucleotide additions commonly observed in T7 RNA polymerase *in vitro* transcription products (*89*). Three peaks are observed in the SLA-NS5 complex spectrum, consistent with SLA:NS5 stoichiometries of 1:1, 2:1 (most prominent) and 3:3.

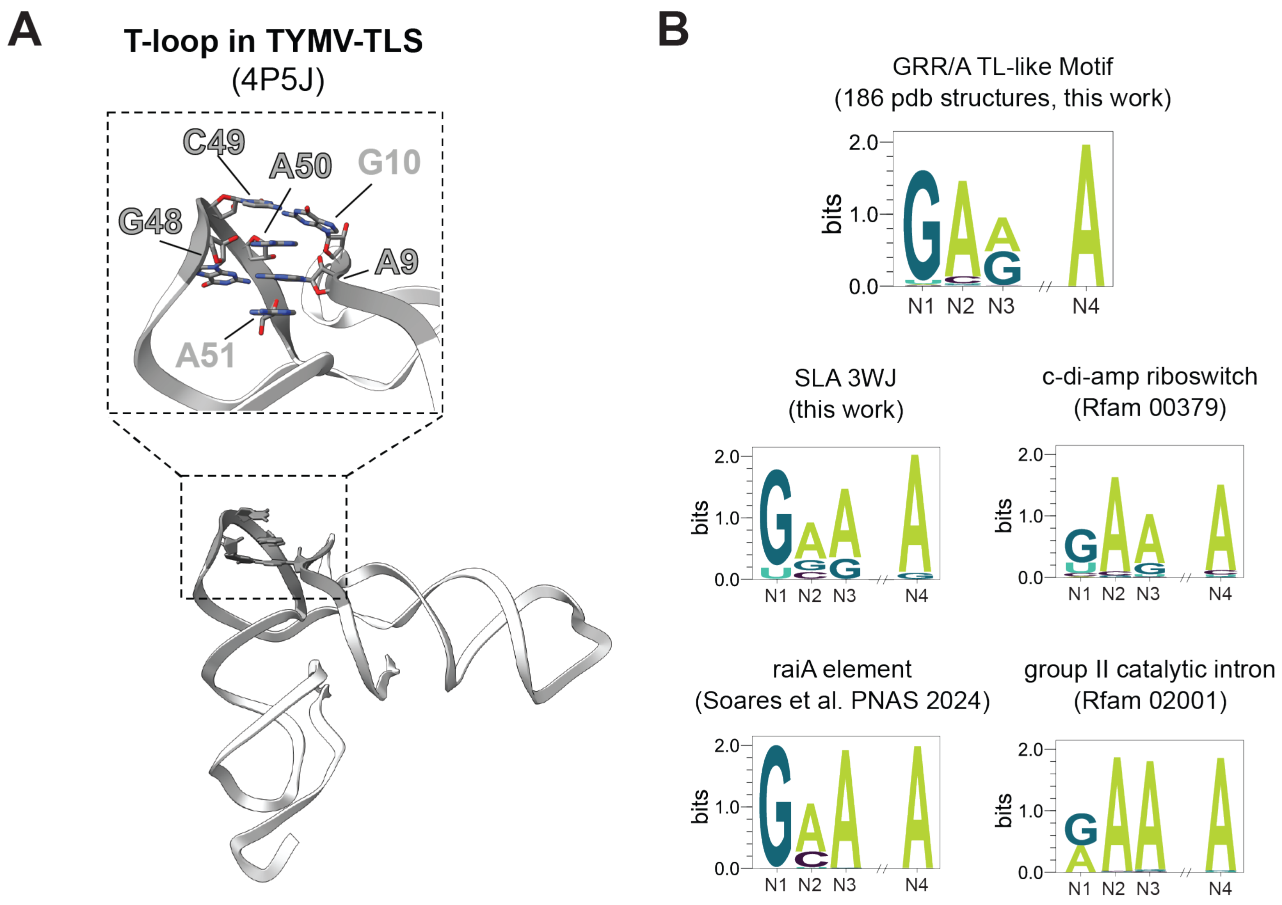

**fig. S20. GRR/A TL-like motif in diverse structural RNAs.** (**A**) GRR/A TL-like motif within a T-loop tertiary interaction in TYMV-TLS (*90*). A9 docks into the T-loop, forming a sheared G:A base-pair with G48 and cross-strand stacking interactions with A50. (**B**) Sequence logos of 3WJs forming the GRR/A TL-like motif in diverse structural RNAs. The GRR/A TL-like motif consensus logo (top) was generated from WebFR3D hits. The RNA-specific sequence logos were generated from covariance model-based alignments using Infernal (*43*).

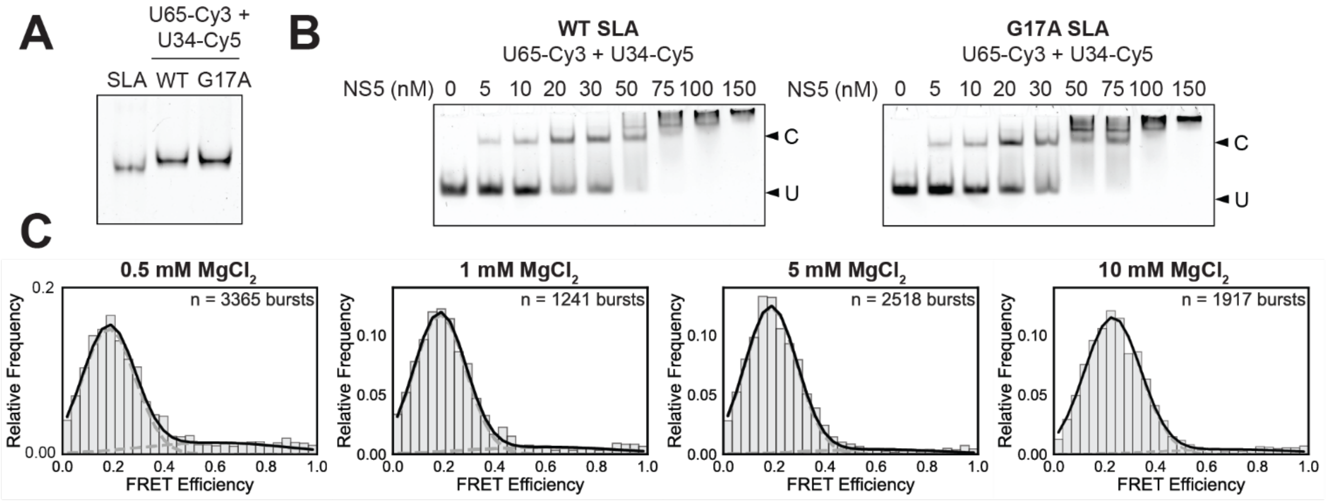

**fig. S21. G17A SLA populates predominantly the S-B state across solution conditions.** (**A**) Native 15% polyacrylamide gel of WT SLA with 3′ end Cy3 label (left), Cy3/Cy5-labeled WT SLA (middle) and Cy3/Cy5-labeled G17A SLA (right). Samples were refolded using the smFRET refolding protocol. (**B**) EMSA of Cy3/Cy5-labeled WT and G17A SLA with NS5, visualized using Cy5 detection. Bands corresponding to unbound SLA (U) and SLA-NS5 complex (C) are labeled. (**C**) Normalized FRET histograms (30 bins) of G17A SLA at varying [$Mg^{2+}$] fitted to two-Gaussian models, confirming strong destabilization of the high FRET state across all solution conditions tested.

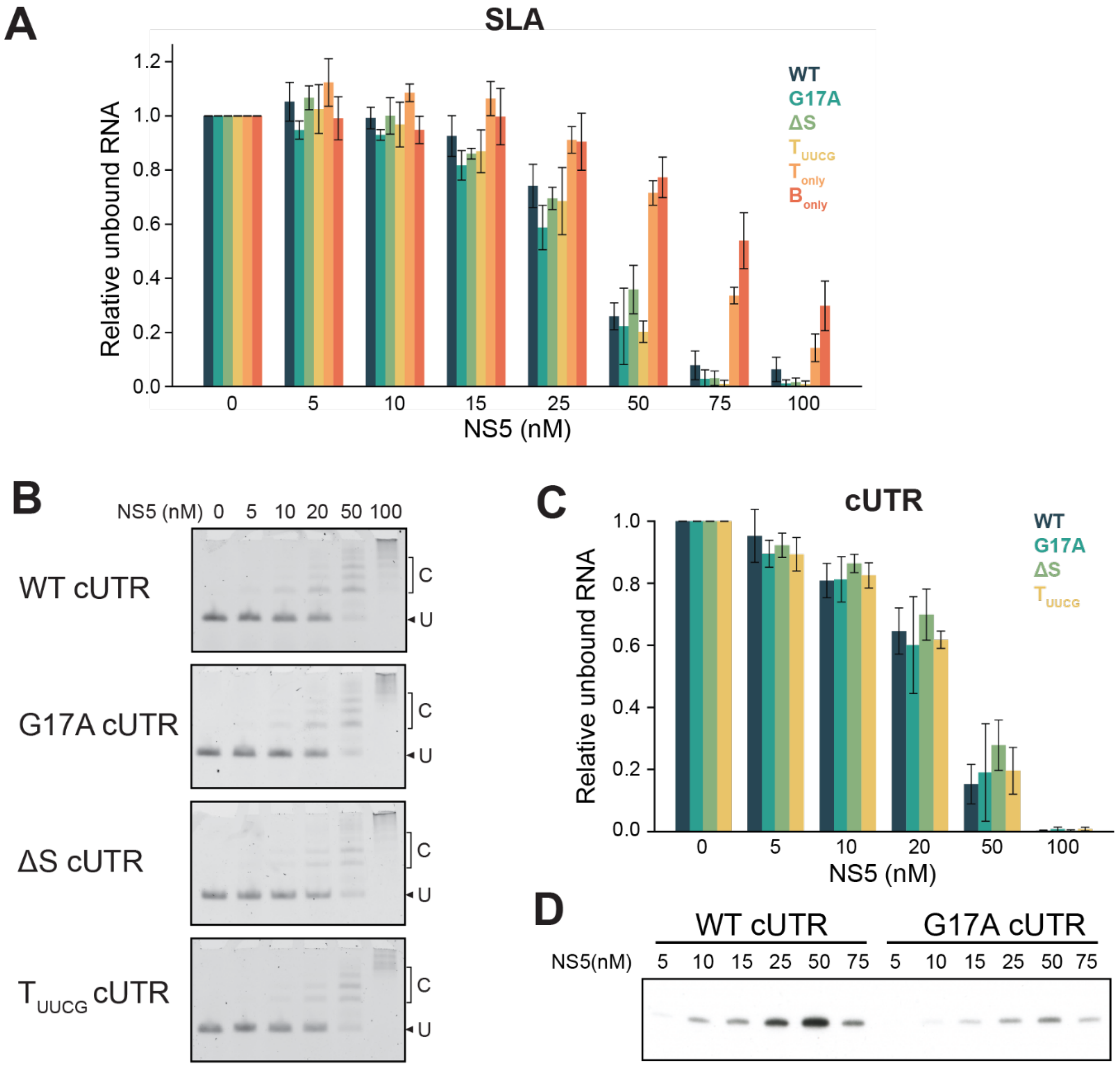

**fig. S22. NS5 binding and negative-strand synthesis assays of DENV2 SLA mutants.** (**A**) Quantification of NS5 binding to SLA variants by EMSA (Fig. 5B,F). Band intensities of unbound RNA were normalized to the band intensity in the absence of NS5. Relative amounts of unbound WT SLA, G17A SLA, ΔS SLA, $T_{UUCG}$ SLA, $T_{only}$ and $B_{only}$ are shown from left to right at each NS5 concentration. (**B**) NS5 binding to cUTR template evaluated by EMSA. Bands corresponding to unbound RNA (U) and NS5-cUTR complex (C) are labeled. (**C**) Quantification of NS5 binding to cUTR from panel B. Band intensities of unbound RNA were normalized to the band intensity in the absence of NS5. Relative amounts of unbound WT, G17A, ΔS and $T_{UUCG}$ cUTRs are shown from left to right at each NS5 concentration. (**D**) NS5 titration for negative-strand synthesis using WT and G17A cUTR templates. Quantification is shown in Fig. 5D.

**Table S1: Predicted inter-dye distances and FRET efficiencies for S-T and S-B states of DENV2 SLA.** Mean inter-dye distances and FRET efficiencies were calculated using accessible volume (AV) simulations in Olga (*91*). Dye positions corresponding to U65 (Cy3) and U34 (Cy5) were used, consistent with the smFRET labeling scheme shown in fig. S8A. AV simulation parameters were as follows: simulation type: AV3, allowed sphere radius: 5, linker length: 21, linker width: 3.5, radius 2: 3, radius 3: 1.5, radius 1 (Cy5): 10, radius 1 (Cy3): 8.

| | **Mean position distance** | **Mean FRET efficiency ($R_0$ = 50 Å)** |
|---|---|---|
| S-B state | 56.7 ± 6.1 Å | 0.34 ± 0.13 |
| S-T state | 36.0 ± 9.6 Å | 0.72 ± 0.17 |

**Table S2: Summary of sequences used.** For gBlocks used for *in vitro* transcription, the T7 promoter sequence is shown in bold and primer annealing sites are underlined. SLA sequences of interest are highlighted in blue. The HDV ribozyme sequence, appended to generate homogeneous 3′ ends, is shown in grey and is removed after *in vitro* transcription upon self-cleavage. Residues bearing internal modifications in custom single-stranded RNA oligos are shown in red. For the NS5 constructs, the N-terminal 6x-His tag and thrombin cleavage site sequences are shown in yellow and green, respectively.

| | |
|---|---|
| DENV2 SLA-TET gBlock | AGTGAATTC**TAATACGACTCACTATA**GGAGGGAAAAGTTATCA<br>GGCATGCACCTGGTAGCTAGTCTTTAAACCAATAGATTGCATC<br>GGTTTAAAAGGCAAGACCGTCAAATTGCGGGAAAGGGGTCAA<br>CAGCCGTTCAGTACCAAGTCTCAGGGGAAACTTTGAGATGGC<br>CTTGCAAAGGGTATGGTAATAAGCTGACGGACATGGTCCTAA<br>CCACGCAGCCAAGTCCTAATTGTTAGTCTACGTGGACCGACA<br>AAGACAGATTCTTTGAGGGAGCTAAGCTCAACGTAGTTCTAAC<br>AGATGGATGCAGTTCACAGACTAAATGTCGGTCGGGGAAGAT<br>GTATTCTTCTCATAAGATATAGTCGGACCTCTCCTTAATGGGA<br>GCTAGCGGATGAAGTGATGCAACACTGGAGCCGCTGGGAACT<br>AATTTGTATGCGAAAGTATATTGATTAGTTTTGGAG |
| DENV2 SLA-cpTET gBlock | AGTGAATTC**TAATACGACTCACTATT**AGTTGTTAGTCTACGTG<br>GACCGACAAAGACAGATTCTTTGAGGGAGCTGATATGGATGC<br>AGTTCACAGACTAAATGTCGGTCGGGGAAGATGTATTCTTCTC<br>ATAAGATATAGTCGGACCTCTCCTTAATGGGAGCTAGCGGAT<br>GAAGTGATGCAACACTGGAGCCGCTGGGAACTAATTTGTATG<br>CGAAAGTATATTGATTAGTTTTGGAGTACTCGTTGGAGGGAAA<br>AGTTATCAGGCATGCACCTGGTAGCTAGTCTTTAAACCAATAG<br>ATTGCATCGGTTTAAAAGGCAAGACCGTCAAATTGCGGGAAA<br>GGGGTCAACAGCCGTTCAGTACCAAGTCTCAGGGGAAACTTT<br>GAGATGGCCTTGCAAAGGGTATGGTAATAAGCTGACGGACAT<br>GGTCCTAACCACGCAGCCAAGTCCTAAGTCAGCTCAACGTAG<br>TTCTAACAG |
| ZIKV SLA-TET gBlock | AGTGAATTC**TAATACGACTCACTATA**GGAGGGAAAAGTTATCA<br>GGCATGCACCTGGTAGCTAGTCTTTAAACCAATAGATTGCATC<br>GGTTTAAAAGGCAAGACCGTCAAATTGCGGGAAAGGGGTCAA<br>CAGCCGTTCAGTACCAAGTCTCAGGGGAAACTTTGAGATGGC<br>CTTGCAAAGGGTATGGTAATAAGCTGACGGACATGGTCCTAA<br>CCACGCAGCCAAGTCCTAATTGTTGATCTGTGTGAGTCAGACT<br>GCGACAGTTCGAGTCTGAAGCGAGAGCTAACAACAGTATCAA<br>CAGATGGATGCAGTTCACAGACTAAATGTCGGTCGGGGAAGA<br>TGTATTCTTCTCATAAGATATAGTCGGACCTCTCCTTAATGGG<br>AGCTAGCGGATGAAGTGATGCAACACTGGAGCCGCTGGGAA<br>CTAATTTGTATGCGAAAGTATATTGATTAGTTTTGGAG |
| WNV SLA-TET gBlock | AGTGAATTC**TAATACGACTCACTATA**GGAGGGAAAAGTTATCA<br>GGCATGCACCTGGTAGCTAGTCTTTAAACCAATAGATTGCATC<br>GGTTTAAAAGGCAAGACCGTCAAATTGCGGGAAAGGGGTCAA<br>CAGCCGTTCAGTACCAAGTCTCAGGGGAAACTTTGAGATGGC |

| | |
|---|---|
| | CTTGCAAAGGGTATGGTAATAAGCTGACGGACATGGTCCTAACCACGCAGCCAAGTCCTAATAGTTCGCCTGTGTGAGCTGACAAACTTAGTAGTGTTTGTGAGGATTAACAACAATTAACACAGTGCGAGCTGATGGATGCAGTTCACAGACTAAATGTCGGTCGGGGAAGATGTATTCTTCTCATAAGATATAGTCGGACCTCTCCTTAATGGGAGCTAGCGGATGAAGTGATGCAACACTGGAGCCGCTGGGAACTAATTTGTATGCGAAAGTATATTGATTAGTTTTGGAG |
| YFV SLA-TET gBlock | AGTGAATTC**TAATACGACTCACTATA**GGAGGGAAAAGTTATCAGGCATGCACCTGGTAGCTAGTCTTTAAACCAATAGATTGCATCGGTTTAAAAGGCAAGACCGTCAAATTGCGGGAAAGGGGTCAACAGCCGTTCAGTACCAAGTCTCAGGGGAAACTTTGAGATGGCCTTGCAAAGGGTATGGTAATAAGCTGACGGACATGGTCCTAACCACGCAGCCAAGTCCTAAAATCCTGTGTGCTAATTGAGGTGCATTGGTCTGCAAATCGAGTTGCTAGGCAATAAACACATTTGGATTATGGATGCAGTTCACAGACTAAATGTCGGTCGGGGAAGATGTATTCTTCTCATAAGATATAGTCGGACCTCTCCTTAATGGGAGCTAGCGGATGAAGTGATGCAACACTGGAGCCGCTGGGAACTAATTTGTATGCGAAAGTATATTGATTAGTTTTGGAG |
| SLA-TET forward primer | AGTGAATTCTAATACGACTCACTATAGG |
| SLA-TET reverse primer | CTCCAAAACTAATCAATATACTTTCGC |
| DENV2 SLA-cpTET forward primer | AGTGAATTCTAATACGACTCACTATTAG |
| DENV2 SLA-cpTET reverse primer | CTGTTAGAACTACGTTGAGCTG |
| DENV2 SLA gBlock | AGTGAATTC**TAATACGACTCACTATT**AGTTGTTAGTCTACGTGGACCGACAAAGACAGATTCTTTGAGGGAGCTAAGCTCAACGTAGTTCTAACAGATGGATGCAGTTCACAGACTAAATGTCGGTCGGGGAAGATGTATTCTTCTCATAAGATATAGTCGGACCTCTCCTTAATGGGAGCTAGCGGATGAAGTGATGCAACACTGGAGCCGCTGGGAACTAATTTGTATGCGAAAGTATATTGATTAGTTTTGGAGTACTCGTTGGAGGGAAAAGTTATCAGGCATGCACCTGGTAGCTAGTCTTTAAACCAATAGATTGCATCGGTTTAAAAGGCAAGACCGTCAAATTGCGGGAAAGGGGTCAACAGCCGTTCAGTACCAAGTCTCAGGGGAAACTTTGAGATGGCCTTGCAAAGGGTATGGTAATAAGCTGACGGACATGGTCCTAACCACGCAGCCAAGTCCTAA |
| DENV2 SLA forward primer | AGTGAATTCTAATACGACTCACTATTAG |
| DENV2 SLA reverse primer | CTGTTAGAACTACGTTGAGCTTAG |
| ZIKV SLA gBlock | AGTGAATTC**TAATACGACTCACTATT**AGTTGTTGATCTGTGTGAGTCAGACTGCGACAGTTCGAGTCTGAAGCGAGAGCTAACAACAGTATCAACAGGGGGCGGCATGGTCCCAGCCTCCTCGCT |
| ZIKV SLA forward primer | AGTGAATTCTAATACGACTCACTATTAG |

| | |
|---|---|
| ZIKV SLA reverse primer | CCTGTTGATACTGTTGTTAGC |
| DENV2 WT cUTR-HDV ribozyme gBlock | GCGAGTGAATTC**TAATACGACTCACTATT**AGTTGTTAGTCTAC<br>GTGGACCGACAAAGACAGATTCTTTGAGGGAGCTAAGCTCAA<br>CGTAGTTCTAACAGTTTTTTAATTAGAGAGCAGATCTCTGTTCG<br>CAGAGATCCTGCTGTCTCCTCAGCATCATTCCAGGCACAGAA<br>CGCCAGAAAATGGAATGGTGCTGTTGAATCAACAGGTTCTGG<br>CCGGCATGGTCCCAGCCTCCTCGCTGGCGCCGGCTGGGCAA<br>CATTCCGAGGGGACCGTCCCCTCGGTAATGGCGAATGGGAC |
| DENV2 G17A cUTR-HDV ribozyme gBlock | GCGAGTGAATTC**TAATACGACTCACTATT**AGTTGTTAGTCTAC<br>GTAGACCGACAAAGACAGATTCTTTGAGGGAGCTAAGCTCAA<br>CGTAGTTCTAACAGTTTTTTAATTAGAGAGCAGATCTCTGTTCG<br>CAGAGATCCTGCTGTCTCCTCAGCATCATTCCAGGCACAGAA<br>CGCCAGAAAATGGAATGGTGCTGTTGAATCAACAGGTTCTGG<br>CCGGCATGGTCCCAGCCTCCTCGCTGGCGCCGGCTGGGCAA<br>CATTCCGAGGGGACCGTCCCCTCGGTAATGGCGAATGGGAC |
| DENV2 ΔS cUTR-HDV ribozyme gBlock | GCGAGTGAATTC**TAATACGACTCACTATT**AGTTGTTAGTCTAC<br>GTGGACCGACAAAGACAGATTCTTTGAGGAACGTAGTTCTAAC<br>AGTTTTTTAATTAGAGAGCAGATCTCTGTTCGCAGAGATCCTG<br>CTGTCTCCTCAGCATCATTCCAGGCACAGAACGCCAGAAAAT<br>GGAATGGTGCTGTTGAATCAACAGGTTCTGGCCGGCATGGTC<br>CCAGCCTCCTCGCTGGCGCCGGCTGGGCAACATTCCGAGGG<br>GACCGTCCCCTCGGTAATGGCGAATGGGAC |
| DENV2 $T_{UUCG}$ cUTR-HDV ribozyme gBlock | GCGAGTGAATTC**TAATACGACTCACTATT**AGTTGTTAGTCTAC<br>GTGGACCGACAAAGATTCGTCTTTGAGGGAGCTAAGCTCAAC<br>GTAGTTCTAACAGTTTTTTAATTAGAGAGCAGATCTCTGTTCGC<br>AGAGATCCTGCTGTCTCCTCAGCATCATTCCAGGCACAGAAC<br>GCCAGAAAATGGAATGGTGCTGTTGAATCAACAGGTTCTGGC<br>CGGCATGGTCCCAGCCTCCTCGCTGGCGCCGGCTGGGCAAC<br>ATTCCGAGGGGACCGTCCCCTCGGTAATGGCGAATGGGAC |
| DENV2 cUTR forward primer | AGTGAATTCTAATACGACTCACTATTAG |
| DENV2 cUTR reverse primer | GTCCCATTCGCCATTAC |
| DENV2 WT SLA smFRET 5′ fragment (containing internal 5-aminohexylacrylamino-uridine modification) | AGUUGUUAGUCUACGUGGACCGACAAAGACAGA**U**UCUUUGA<br>G |
| DENV2 G17A SLA smFRET 5′ fragment (containing internal 5-aminohexylacrylamino-uridine modification) | AGUUGUUAGUCUACGUAGACCGACAAAGACAGA**U**UCUUUGA<br>G |
| DENV2 WT/G17A SLA smFRET 3′ fragment | GGAGCUAAGCUCAACGUAGUUC**U**AACAG |

| | |
|---|---|
| (containing internal 5-aminohexylacrylamino-uridine modification and 5′-phosphate) | |
| SLA smFRET DNA splint for T4 RNA ligase | GTTGAGCTTAGCTCCCTCAAAGAATCTGTC |
| DENV2 WT SLA 3′-Cy3 RNA | AGUUGUUAGUCUACGUGGACCGACAAAGACAGAUUCUUUGA GGGAGCUAAGCUCAACGUAGUUCUAACAGUUU |
| DENV2 G17A SLA 3′-Cy3 RNA | AGUUGUUAGUCUACGUAGACCGACAAAGACAGAUUCUUUGA GGGAGCUAAGCUCAACGUAGUUCUAACAGUUU |
| DENV2 ΔS SLA 3′-Cy3 RNA | AGUUGUUAGUCUACGUGGACCGACAAAGACAGAUUCUUUGA GGAACGUAGUUCUAACAGUUU |
| DENV2 $T_{UUCG}$ SLA 3′-Cy3 RNA | AGUUGUUAGUCUACGUGGACCGACAAAGAUUCGUCUUUGAG GGAGCUAAGCUCAACGUAGUUCUAACAGUUU |
| DENV2 $B_{only}$ 3′-Cy3 RNA | AGUUGUUAGUCUACGUUUCGACGUAGUUCUAACAGUUU |
| DENV2 $T_{only}$ 3′-Cy3 RNA | CCGACAAAGACAGAUUCUUUGAGGUUCG |
| DENV2 NS5 | MGSSHHHHHHSSGLVPRGSHMMKNTANTRRGTGNTGETLGEK WKNRLNALGKSEFQIYKKSGIQEVDRTLAKEGIKRGETDHHAVS RGSAKLRWFVERNLVTPEGKVVDLGCGRGGWSYYCGGLKNVK EVKGLTKGGPGHEEPIPMSTYGWNLVRLQSGVDVFFTPPEKCD TLLCDIGESSPNPTVEAGRTLRVLNLVENWLNNNTQFCIKVLNPY MPSVIEKMEALQRKYGGALVRNPLSRNSTHEMYWVSNASGNIV SSVNMISRMLINRFTMRHKKATYEPDVDLGSGTRNIGIESETPNL DIIGKRIEKIKQEHETSWHYDQDHPYKTWAYHGSYETKQTGSAS SMVNGVVRLLTKPWDVIPMVTQMAMTDTTPFGQQRVFKEKVDT RTQEPKEGTKKLMKITAEWLWKELGKKKTPRMCTREEFTRKVRS NAALGAIFTDENKWKSAREAVEDSGFWELVDKERNLHLEGKCET CVYNMMGKREKKLGEFGKAKGSRAIWYMWLGARFLEFEALGFL NEDHWFSRENSLSGVEGEGLHKLGYILRDVSKKEGGAMYADDT AGWDTRITLEDLKNEEMVTNHMEGEHKKLAEAIFKLTYQNKVVR VQRPTPRGTVMDIISRRDQRGSGQVVTYGLNTFTNMEAQLIRQM EGEGVFKSIQQLTATEEIAVKNWLVRVGRERLSRMAISGDDCVV KPLDDRFASALTALNDMGKVRKDIQQWEPSRGWNDWTQVPFC SHHFHELIMKDGRVLVVPCRNQDELIGRARISQGAGWSLRETAC LGKSYAQMWSLMYFHRRDLRLAANAICSAVPSHWVPTSRTTWSI HATHEWMTTEDMLTVWNRVWIQENPWMEDKTPVESWEEIPYL GKREDQWCGSLIGLTSRATWAKNIQTAINQVRSLIGNEEYTDYM PSMKRFRREEEEAGVLW |
| ZIKV NS5 | MGSSHHHHHHSSGLVPRGSHGGGTGETLGEKWKARLNQMSAL EFYSYKKSGITEVCREEARRALKDGVATGGHAVSRGSAKLRWLV ERGYLQPYGKVIDLGCGRGGWSYYAATIRKVQEVKGYTKGGPG HEEPVLVQSYGWNIVRLKSGVDVFHMAAEPCDTLLCDIGESSSS |

| | |
|---|---|
| | PEVEEARTLRVLSMVGDWLEKRPGAFCIKVLCPYTSTMMETLER LQRRYGGGLVRVPLSRNSTHEMYWVSGAKSNTIKSVSTTSQLLL GRMDGPRRPVKYEEDVNLGSGTRAVVSCAEAPNMKIIGNRIERI RSEHAETWFFDENHPYRTWAYHGSYEAPTQGSASSLINGVVRL LSKPWDVVTGVTGIAMTDTTPYGQQRVFKEKVDTRVPDPQEGT RQVMSMVSSWLWKELGKHKRPRVCTKEEFINKVRSNAALGAIFE EEKEWKTAVEAVNDPRFWALVDKEREHHLRGECQSCVYNMMG KREKKQGEFGKAKGSRAIWYMWLGARFLEFEALGFLNEDHWM GRENSGGGVEGLGLQRLGYVLEEMSRIPGGRMYADDTAGWDT RISRFDLENEALITNQMEKGHRALALAIIKYTYQNKVVKVLRPAEK GKTVMDIISRQDQRGSGQVVTYALNTFTNLVVQLIRNMEAEEVLE MQDLWLLRRSEKVTNWLQSNGWDRLKRMAVSGDDCVVKPIDD RFAHALRFLNDMGKVRKDTQEWKPSTGWDNWEEVPFCSHHFN KLHLKDGRSIVVPCRHQDELIGRARVSPGAGWSIRETACLAKSYA QMWQLLYFHRRDLRLMANAICSSVPVDWVPTGRTTWSIHGKGE WMTTEDMLVVWNRVWIEENDHMEDKTPVTKWTDIPYLGKREDL WCGSLIGHRPRTTWAENIKNTVNMVRRIIGDEEKYMDYLSTQVR YLGEEGSTPGVL |

**Table S3: Constraints matrix input to WebFR3D to search for GRR/A TL-like motif.** Positions 1-3 are enforced to be consecutive, positions 3 and 4 must be separated by more than 3 nucleotides, positions 1 and 4 must form sugar-Hoogsteen base-pair, and positions 2,3 and 4 must form stacking interactions. This search returned 3227 hits, which were subsequently filtered to remove duplicate entries referring to the same RNA deposited multiple times in the PDB, resulting in the final 186 unique hits.

| | **Position 1** | **Position 2** | **Position 3** | **Position 4** |
|---|---|---|---|---|
| **Position 1** | | | ~pair | tSH |
| **Position 2** | =1 > | | stack and ~s55 | ~pair |
| **Position 3** | | =1 > | | stack and ~s55 |
| **Position 4** | | | => 3 | |

**Table S4: Summary of cryo-EM sample preparation and data collection parameters.** The final sample concentrations, buffer conditions, grid type, pixel size, exposure time, total dose and spherical aberration are listed for each dataset. *Note: For the ZIKV SLA-NS5 dataset, approximately 20% of micrographs were collected at a 20° tilt.

| Name | Concentration | Buffer | Grid type | Pixel size (Å) | Exposure time (s) | Total dose (e/Å$^2$) | Spherical aberration |
|---|---|---|---|---|---|---|---|
| DENV2 SLA | 75 μM | 50 mM Tris-HCl pH 7.5, 10 mM $MgCl_2$ | C-Flat CF-1.2/1.3-4Cu-T-50 400 mesh | 0.2575 | 1.2 | 54.3 | 0 |
| DENV2 SLA-TET | 20 μM | 50 mM Tris-HCl pH 7.5, 10 mM $MgCl_2$ | C-Flat CF-1.2/1.3-4Cu-T-50 400 mesh | 0.2575 | 1.3 | 57.9 | 0 |
| DENV2 SLA-cpTET | 20 μM | 50 mM Tris-HCl pH 7.5, 10 mM $MgCl_2$ | C-Flat CF-1.2/1.3-4Cu-T-50 400 mesh | 0.2575 | 1.3 | 59.7 | 0 |
| ZIKV SLA-TET | 20 μM | 50 mM Tris-HCl pH 7.5, 10 mM $MgCl_2$ | C-Flat CF-1.2/1.3-4Cu-T-50 400 mesh | 0.2686 | 1.6 | 60.6 | 2.7 |
| WNV SLA-TET | 20 μM | 50 mM Tris-HCl pH 7.5, 10 mM $MgCl_2$ | C-Flat CF-1.2/1.3-4Cu-T-50 400 mesh | 0.264 | 1.4 | 60.4 | 0 |
| YFV SLA-TET | 20 μM | 50 mM Tris-HCl pH 7.5, 10 mM $MgCl_2$ | C-Flat CF-1.2/1.3-4Cu-T-50 400 mesh | 0.2575 | 1.2 | 54.3 | 0 |
| DENV2 SLA-NS5 | 2.5 μM SLA + 2.5 μM NS5 | 50 mM Tris pH 7.5, 75 mM NaCl, 1 mM $MgCl_2$, 5 mM DTT, 5% glycerol | C-Flat CF-1.2/1.3-4Cu-T-50 400 mesh | 0.2575 | 1.2 | 54.3 | 0 |
| ZIKV SLA-NS5* | 2 μM SLA + 2 μM NS5 | 50 mM Tris-HCl pH 7.5, 75 mM NaCl, 1 mM $MgCl_2$, | QuantiFoil, R2/1, 200 mesh, Cu H2 | 0.515 | 1.2 | 54.3 | 0 |

| | | |
|---|---|---|
| | | 5 mM DTT, 2.5% glycerol |
